# OCellus: A Language-Model Framework for Single-Cell, Spatial, and Perturbation Biology with Natural-Language Reasoning

**DOI:** 10.64898/2026.07.08.737248

**Authors:** Chen Zhang, Jianlong Sun, Zaoxu Xu, Renjie Liao, An Yin, Hui Gao, Erkai Liu, Yuanye Bao, Luyang Zhao, Gufeng Wang

## Abstract

Computational modeling of cellular behavior—the virtual cell—has emerged as a stated grand challenge at the intersection of artificial intelligence and biology, yet existing foundation models remain specialized: single-cell models process dissociated transcriptomes only, spatial models require dedicated spatial-aware architectures, and perturbation predictors depend on manually curated knowledge bases that cap generalization. Here we introduce OCellus, a single nine-billion-parameter language model (Qwen3.5-9B) fine-tuned on twenty-two biological tasks that simultaneously addresses all three limitations through three coordinated technical contributions on a shared backbone. First, EvenClock encodes two-dimensional spatial coordinates as eighteen clockface sectors of text, enabling spatial reasoning on a vanilla language model without architectural modification; on ten spatial transcriptomics tasks OCellus attains 77 percent spatial-neighborhood accuracy, 96 percent spatial-cellchat accuracy, and 0.70 proportion-cosine similarity on spatial deconvolution, all without any spatial-aware architectural components. Second, per-gene language-model embeddings replace the Gene Ontology annotations that GEARS depends on, achieving Pearson correlation 0.945 on the Replogle 2022 perturbation benchmark versus 0.84 for GEARS across 457 completely unseen knockout genes. Third, OCellus-Agent provides a Planner–Router–Verifier natural-language interface that achieves 75 percent pipeline accuracy on eighty multi-task queries. Removing language-model embeddings collapses perturbation Pearson to 0.06, confirming that learned functional representations—not graph topology—drive the gain. As a cell-type encoder, OCellus ranks first among fourteen foundation models in linear-probe accuracy at 95.1 percent across four benchmark datasets, and reaches 72.6 percent average across twenty-two evaluated biological tasks—a 57-percentage-point absolute gain over the strongest baseline configuration. As a language model, OCellus uniquely generates natural-language explanations of its predictions, a capability absent from all competing methods. Code, pre-trained model weights, the graph-neural-network module, and the agent system will be made available upon publication.

## Introduction

Computational modeling of cellular behavior—the virtual cell—has emerged as a grand challenge at the intersection of artificial intelligence and biology (Bunne et al., 2024). A comprehensive virtual cell would predict how a genetic perturbation reshapes the transcriptome, classify cell identity from a gene-expression profile, reconstruct tissue architecture from spatially resolved measurements, and explain its predictions in biologically interpretable language. Realizing this vision would compress decades of accumulated biological knowledge into an executable form, with high application value for in silico perturbation screening, cell-type atlas construction, spatial pathology, and clinical decision support.

Recent advances in foundation models have brought this vision closer to reality. Transformer-based architectures pretrained on tens of millions of cellular profiles—scGPT (Cui et al., 2024), Geneformer (Theodoris et al., 2023), scFoundation (Hao et al., 2024), UCE (Rosen et al., 2023), and CellPLM (Wen et al., 2024)—yield transferable representations of gene programs and cell states. For spatial transcriptomics, SToFM, Nicheformer (2025), and HEIST (2025) extend representation learning to tissue context but require dedicated spatial-aware architectures; concurrent work applying language models directly to single-cell or spatial data includes CeLLM (Li et al., 2024), ST-LLM (Wang et al., 2024), Cell2Sentence (C2S, Ou et al., 2024), C2S-Scale (Yang et al., 2024), and AlphaCell (2025), which we discuss in turn below. In perturbation prediction, GEARS (Roohani et al., 2024) established graph neural networks over Gene Ontology as the academic benchmark, achieving Pearson correlation 0.84 on the Norman 2019 dataset.

Despite this rapid progress, three classes of specialized foundation models have emerged, each advancing one capability but none addressing all. These limitations collectively prevent any existing foundation model from serving as a true virtual cell.

The first limitation is modality fragmentation. Single-cell foundation models such as scGPT, Geneformer, and scFoundation process dissociated RNA-seq profiles and produce strong cell-state representations but cannot directly incorporate spatial context. Spatial foundation models such as SToFM and HEIST extend representation learning to tissue context, yet they require dedicated spatial-aware architectures—hierarchical graph transformers in HEIST and SE(2)-equivariant networks in SToFM— that cannot share parameters with single-cell counterparts. Consequently, a model trained on millions of dissociated cells cannot directly inform spatial tissue analysis, and spatial models lack the broad cell-type coverage of single-cell atlases. Nicheformer partially bridges this gap by jointly training spatial and dissociated data within a unified transformer. As a result, currently there is no existing foundation model that simultaneously unifies dissociated single-cell, spatial transcriptomics, and perturbation prediction within a single, unmodified backbone.

The second limitation is curated-knowledge dependence in perturbation generalization. The standard Replogle 2022 benchmark (Replogle et al., 2022) requires predicting transcriptional responses for 457 knockout genes that share zero overlap with the 1,830 genes observed during training. Under these conditions, even GEARS achieves direction accuracy of only 54.6 percent—barely exceeding the 50 percent random baseline—because Gene Ontology annotations are incomplete for many genes and do not capture the full complexity of regulatory relationships. The fundamental problem is that models must generalize perturbation effects to genes whose functional profiles are only partially characterized.

The third limitation is opacity. All current virtual-cell models operate as black-box predictors: they output numerical predictions without biological reasoning. A researcher using GEARS learns that knocking out TP53 changes MDM2 expression by a log2 fold-change of −1.8, but the model cannot explain why. This opacity limits scientific discovery, clinical trust, and the model’s utility as a hypothesis-generation tool.

Here we present OCellus, a single nine-billion-parameter language model that addresses all three limitations through three coordinated technical contributions on a shared Qwen3.5-9B backbone (Bai et al., 2025). The model is fine-tuned via LoRA (Hu et al., 2022) on twenty-two biological tasks spanning dissociated single-cell and spatially resolved transcriptomics, including novel tasks such as cross-species cell-type annotation, developmental-stage prediction, and binary protein–protein interaction classification on the STRING database (Szklarczyk et al., 2023). First, EvenClock discretizes the continuous two-dimensional plane into a clockface of six directions and three distance rings, encoding each sector’s neighboring cells as a gene-list prefix that the language model consumes as text with no architectural modification. Second, per-gene language-model embeddings extracted from the frozen OCellus backbone replace the Gene Ontology annotations that GEARS depends on, serving as node features for a plug-in graph neural network over the STRING v12 protein-interaction graph. Third, OCellus-Agent organizes the full capability suite as a Planner–Router–Verifier system that accepts natural-language biological queries and produces executable directed acyclic graphs of typed tool calls, with a three-layer Verifier providing biological plausibility checks. The unifying insight is that multi-task language-model training yields a functional representation of every gene token that simultaneously drives spatial reasoning, perturbation generalization, and natural-language explanation; the full architecture is detailed in Results §2.1. Empirically, OCellus attains Pearson correlation 0.945 on Replogle 2022 perturbation prediction, the OCellus encoder ranks first among fourteen foundation models in linear-probe accuracy at 95.1 percent, and OCellus-Agent reaches 75 percent pipeline accuracy on natural-language biological queries.

## Results

### The OCellus framework

OCellus is organized around three components that share a single Qwen3.5-9B backbone: a multi-task language model that serves simultaneously as encoder, predictor, and explainer; a plug-in graph neural network that handles continuous perturbation prediction; and an agent layer that exposes the full capability suite through natural language (Figure 1). The same nine-billion-parameter model thus produces cell-type embeddings ranked first among fourteen foundation models, executes twenty-two biological tasks at expert-level accuracy, predicts transcriptional responses to unseen genetic knockouts with state-of-the-art fidelity, and generates natural-language explanations of its own outputs. The three roles—encoder, predictor, explainer—are co-located on a shared backbone via dynamic adapter swapping. This arrangement distinguishes OCellus from prior foundation models that populate at most one or two of these roles with separate models; the roles share the same backbone weights but are activated by separate LoRA adapters trained on different task mixes.

**Figure 1.**
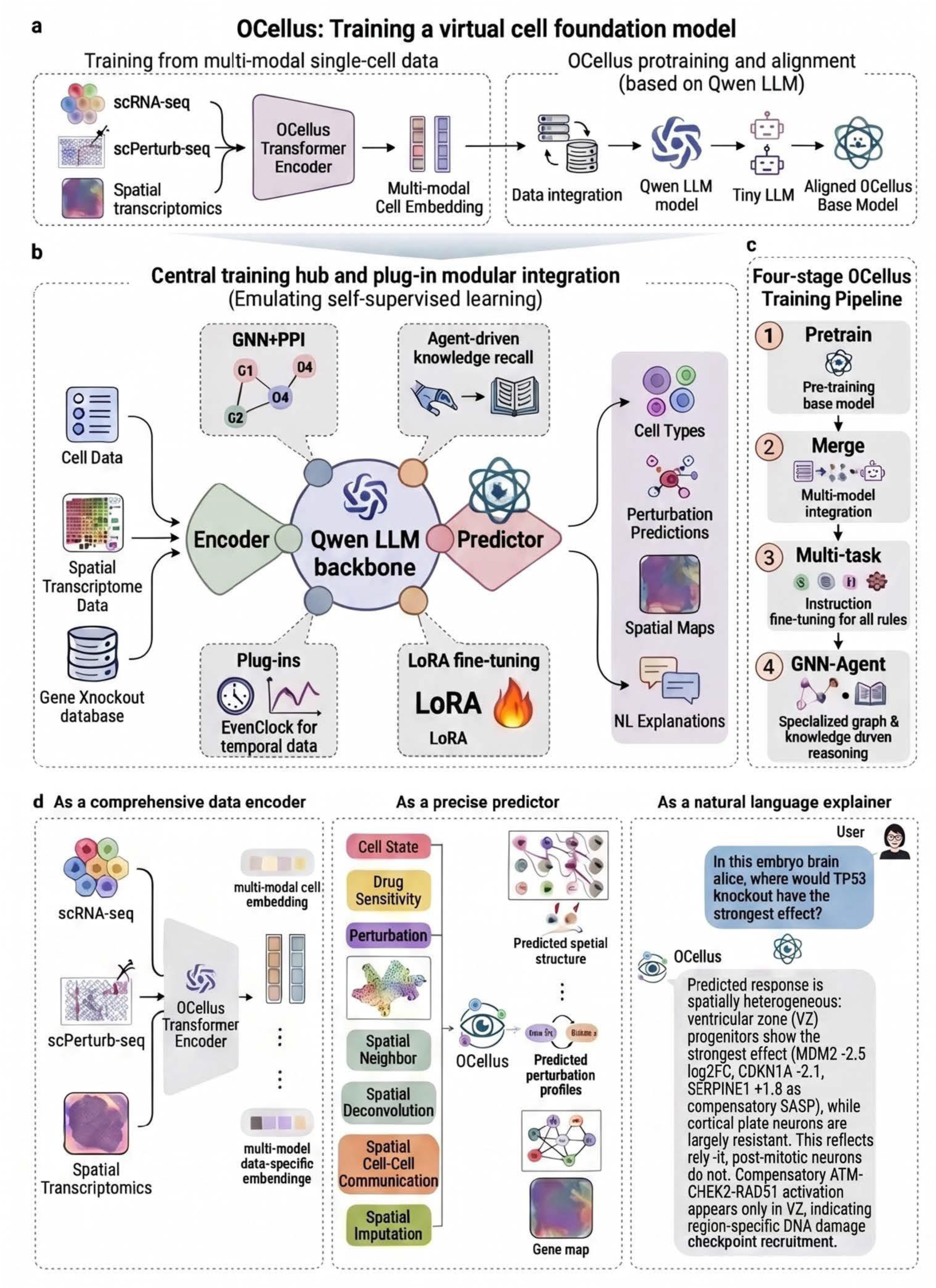
OCellus framework overview (architectural schematic, not empirical data). (a) Training overview: multi-modal biological data (single-cell RNA-seq, spatial transcriptomics, perturbation screens, protein-protein interaction databases) is tokenized and passed through a transformer encoder, then aligned to a Qwen large-language-model backbone to produce the OCellus base model. (b) Central training hub and plug-in modular integration: the Qwen LLM backbone sits at the center, with attached plug-in components including the graph neural network over protein-protein interaction graphs, the agent knowledge-recall module, the EvenClock spatial-encoding module, and the LoRA adapters that activate per task. (c) Four-stage OCellus training pipeline: pretraining on representation-learning tasks, multi-model adapter merging to produce the OCellus-Pretrain checkpoint, multi-task instruction tuning to produce the OCellus checkpoint, and final graph-neural-network and agent LoRA training. (d) Three functional roles served by the same backbone: encoder (multi-modal cell and gene embeddings), predictor (cell-state, perturbation, and spatial tasks), and explainer (natural-language question answering and biological reasoning).

The backbone is Qwen3.5-9B, fine-tuned via LoRA with rank sixty-four, scaling factor sixty-four, rank-stabilized LoRA enabled, dropout 0.05, and all attention and MLP projections targeted. We do not unfreeze the embedding or language-modeling-head layers; the rationale and the underlying ablation are documented in Methods (Video-memory optimization). A defining capability of the resulting model is the generation of fixed-length 4,096-dimensional gene embeddings obtained by passing individual gene symbols through the model and averaging last-layer hidden states. These embeddings are cell-type-independent—they encode what each gene does rather than where it is expressed—and serve as the functional-knowledge substrate that drives perturbation generalization, spatial reasoning, and natural-language explanation.

For perturbation prediction, a lightweight graph neural network of approximately 0.6 million parameters operates on the STRING v12 protein–protein interaction network (15,309 nodes, 833,196 edges at confidence ≥ 0.70). Per-gene language-model embeddings from the frozen OCellus backbone serve as initial node features; binary indicators mark the knockout gene and baseline-expression genes; three graph-convolution layers propagate this signal through the network; and a multi-layer-perceptron head produces continuous log2 fold-change predictions. The graph neural network adds less than one million trainable parameters—three orders of magnitude smaller than the nine-billion-parameter backbone—and trains in hours on a single GPU. The full specification appears in Methods.

The agent layer addresses a complementary limitation: even a powerful model is accessible only by writing custom code per task (loading h5ad, applying LoRA, decoding predictions, plotting results). OCellus-Agent organizes the full capability suite as a Coordinator–Expert–Verifier system over the same Qwen3.5-9B backbone. The Coordinator contains a trained Planner LoRA that decomposes natural-language queries into executable directed acyclic graphs of typed tool calls, a rule-based Router that dispatches each step to the appropriate expert role, and a three-layer Verifier that provides biological plausibility checks. Five expert roles—Data Loader, Annotator, Perturbation, Spatial, and Explainer—are implemented as dynamic LoRA swaps on the shared backbone, and a Tool Registry of fifteen registered tools is exposed via a Gradio web interface. Six interchangeable runtime configurations are loaded on demand through dynamic adapter swapping, so the entire agent occupies approximately twenty-two gigabytes of GPU memory on a single GPU rather than five separate model instances. For spatial transcriptomics, EvenClock discretizes two-dimensional coordinates into eighteen clockface sectors of text, allowing the same backbone to consume spatial context with no architectural modification; the spatial-encoding scheme is detailed in Results §2.4.

Complete hyperparameters for all components (LLM backbone, training pipeline, graph neural network, agent LoRAs) are tabulated in Supplementary Table S8. Training proceeds through a four-stage pipeline: pretraining on seven representation-learning tasks, adapter merging to produce the OCellus-Pretrain checkpoint, multi-task fine-tuning on twenty-two tasks, and a final merge to produce the OCellus checkpoint used throughout this paper. Total wall time is 35.5 hours on four NVIDIA A800-80GB GPUs, with a final training loss of 0.6649. The full training specification, including GPU-memory optimization and data preprocessing decisions, appears in Methods and Supplementary Figure S1.

### Multi-task competence across twenty-two biological tasks

We first asked whether multi-task fine-tuning on twenty-two biological tasks confers capability that is absent from base language models. OCellus was evaluated on twelve single-cell tasks—spanning perturbation prediction, gene-interaction classification (synthetic lethality and phenotypic interaction), gene–disease association, cell-marker identification, drug sensitivity, perturbation-response gene ranking, cross-species cell-type annotation, developmental-stage prediction across mouse (E9.5–E16.5) and human (CS12–CS23) embryogenesis, binary protein–protein interaction classification on STRING v12, and drug-treatment cell-state transition prediction—and ten spatial transcriptomics tasks including cell-type annotation, tissue-region segmentation, cell–cell communication inference, neighborhood prediction, gene imputation, density classification, deconvolution, spatially variable gene detection, spatial-domain identification, and cell-type-specific marker prediction. Representative predictions across six task categories are shown in Table 1.

**Table 1.**
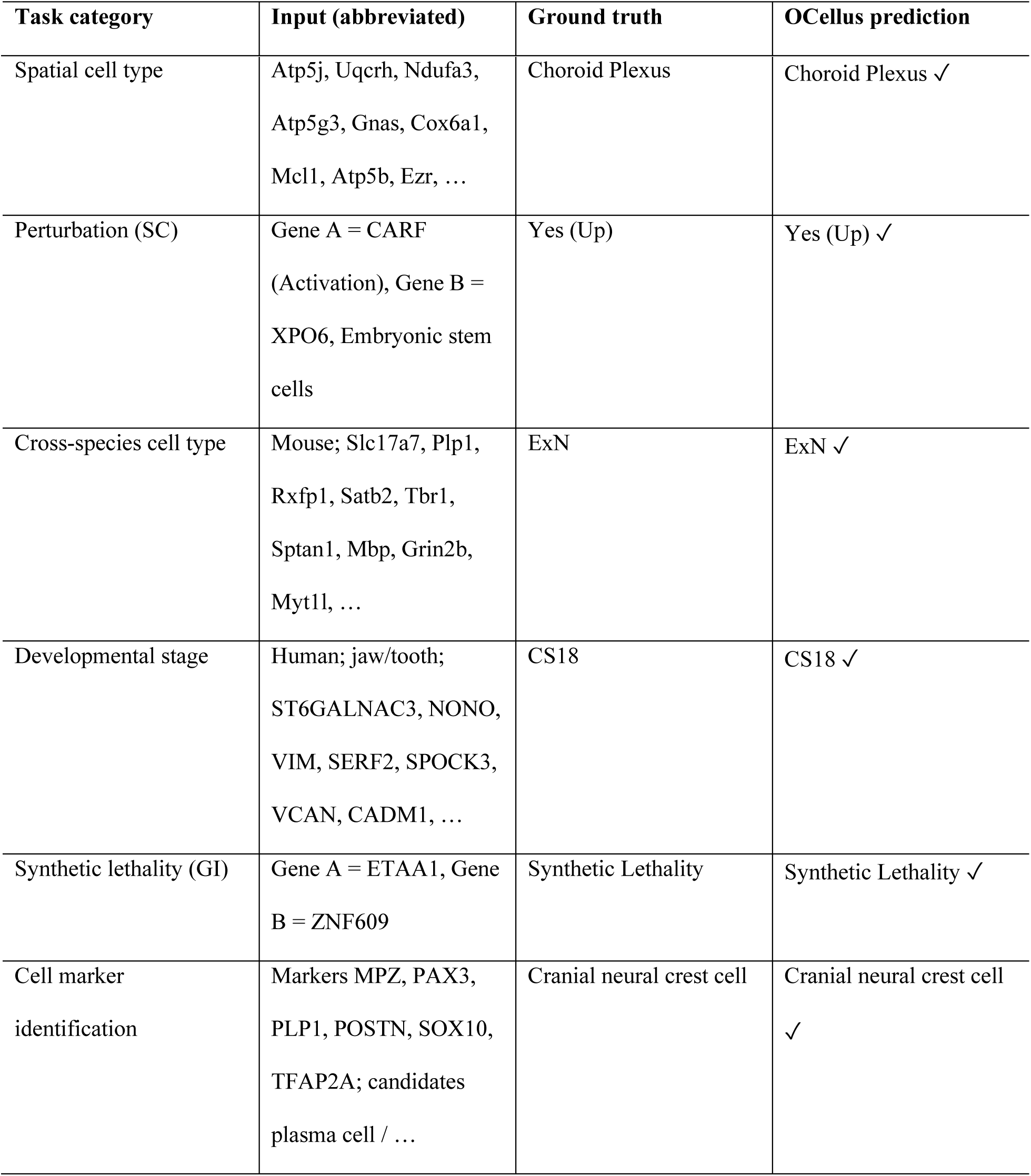
Representative OCellus predictions across six task categories (all six predictions match ground truth; inputs abbreviated for display).

To isolate the contribution of OCellus’s training pipeline, we benchmarked three configurations of the same Qwen3.5-9B backbone under identical protocols: the base Qwen3.5-9B model, OCellus-Pretrain (the model state after Stage 2 pretraining merge but before multi-task fine-tuning), and the full OCellus model (Figure 2a). The primary metric is balanced accuracy—the mean of per-class recall—for short-answer classification tasks, which penalizes models that exploit class imbalance such as a language model that always outputs the majority-class label; for ranking and generation tasks we use gene-set recall against the curated ground-truth gene list; and for spatial deconvolution we use proportion-vector cosine similarity. Full metric definitions appear in Methods.

**Figure 2.**
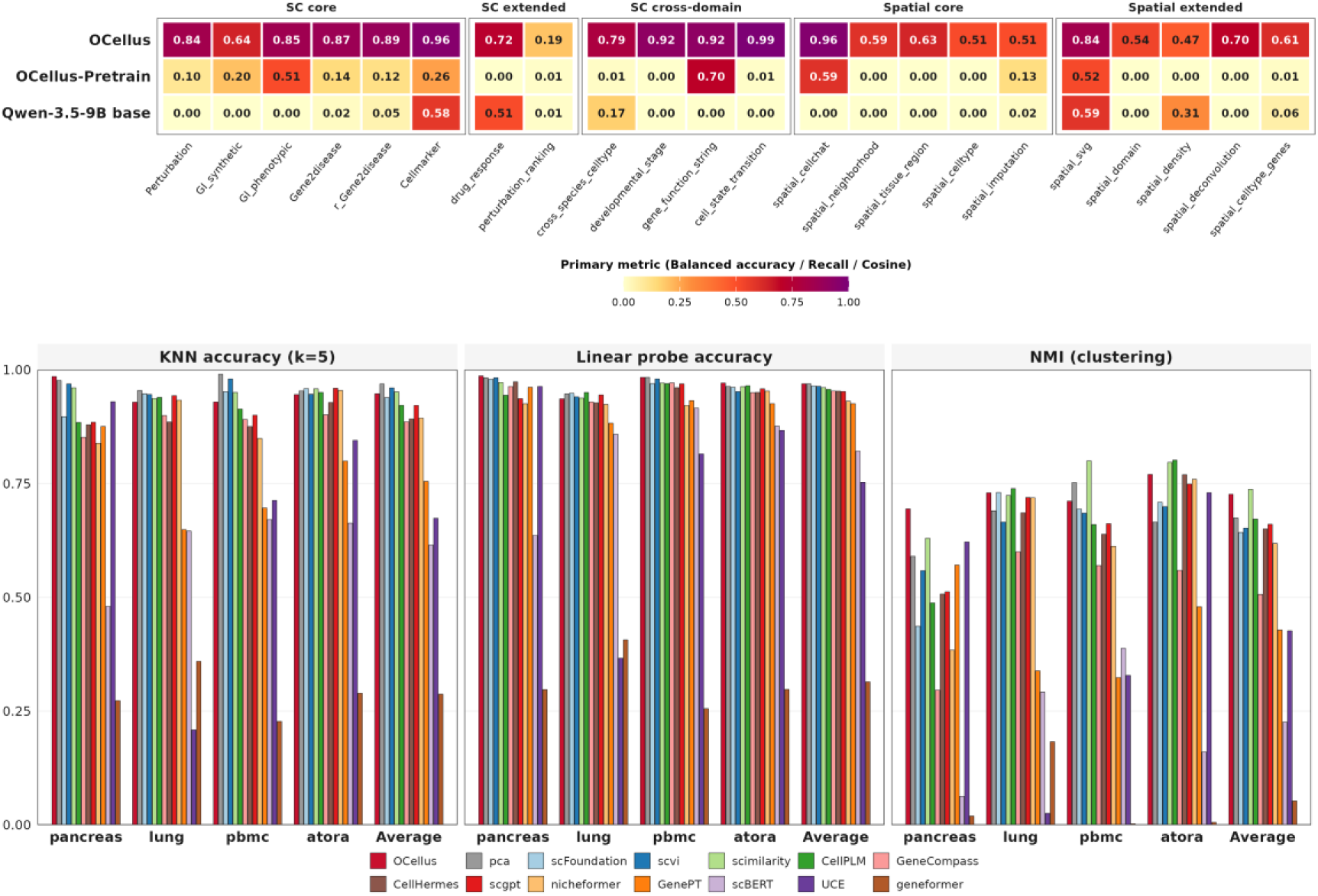
Multi-task benchmark and encoder quality. (a) Multi-task heatmap across 22 task types and 3 model configurations of the same Qwen3.5-9B backbone: base model, OCellus-Pretrain (after Stage 2 merging), and the full OCellus model — all evaluated under identical protocols on held-out test sets. The color scale encodes the task-appropriate primary metric: balanced accuracy for short-answer classification tasks, gene-set recall for ranking/generation tasks, and proportion-vector cosine similarity for spatial deconvolution. Tasks are grouped by category on the x-axis: single-cell core (6), single-cell extended (2), single-cell cross-domain (4), spatial core (5), and spatial extended (5). The full OCellus model achieves 72.6% average — a 57-percentage-point absolute gain over the strongest baseline (OCellus-Pretrain), with the largest improvements on cross-species, developmental-stage, and gene-function tasks that prior cell-language models could not learn. (b) Encoder benchmark across 14 foundation models: kNN accuracy (k=5), linear probe accuracy, and NMI (clustering) across four datasets — pancreas, lung, PBMC, aorta — with an Average summary group. OCellus achieves the highest linear-probe and k-nearest-neighbor accuracy among the fourteen benchmarked models.

The base Qwen3.5-9B model scores 10.5 percent average across the twenty-two evaluated tasks after stripping Qwen’s reasoning prefix that would otherwise prevent short-label parsing. OCellus-Pretrain improves only marginally to 15.1 percent, indicating that pretraining alone does not confer multi-task competence. The full OCellus model reaches 72.6 percent average—a 57-percentage-point absolute gain over the strongest baseline configuration. The largest gains appear on the cross-domain tasks that no prior cell-language model has been trained on, including cross-species cell-type annotation (79.1 percent balanced accuracy), developmental-stage prediction (92.2 percent), STRING protein–protein interaction classification (92.4 percent), and drug-treatment cell-state transition (99 percent gene-set recall). A full per-task heatmap is shown in Figure 2a; complete numbers for all six metrics (balanced accuracy, exact match, gene Jaccard, gene recall, gene precision, proportion cosine) across all twenty-two tasks are reported in Supplementary Table S6 and visualized in Supplementary Figure S2.

We reconcile the 72.6 percent multi-task average with the 95.1 percent linear-probe encoder accuracy reported in the next subsection as follows. The encoder benchmark is a closed-set classification on a moderate number of cell types evaluated on held-out cells from the same datasets used for embedding extraction, whereas the multi-task benchmark covers twenty-two distinct task types including ranking, generation, and tasks with hundreds of labels (for example, the cell-marker identification task alone has 295 classes). The twenty-three-percentage-point gap therefore reflects task-difficulty heterogeneity rather than encoder weakness.

### Best-in-class cell representations

Beyond multi-task performance, we asked whether OCellus produces competitive cell-type embeddings against existing foundation models. We evaluated OCellus against thirteen competing foundation models—including scGPT (Cui et al., 2024), Geneformer (Theodoris et al., 2023), scFoundation (Hao et al., 2024), UCE (Rosen et al., 2023), CellPLM (Wen et al., 2024), Nicheformer (2025), scVI (Lopez et al., 2018), and PCA—using linear-probe and k-nearest-neighbor classification across four standard datasets totaling 73,655 cells and 52 cell types (Figure 2b). In linear-probe classification, OCellus ranks first among all fourteen models with an average accuracy of 95.1 percent (complete per-dataset numbers for all fourteen models are reported in Supplementary Table S7 and visualized in Supplementary Figure S3), surpassing PCA (93.5 percent), CellPLM (94.4 percent), and scGPT (88.4 percent). In k-nearest-neighbor classification at k equal to five, OCellus again ranks first. In unsupervised clustering quality measured by normalized mutual information, OCellus ranks second (NMI = 0.71) behind only scimilarity, a tool specifically optimized for cell-type similarity search.

UMAP visualization of pancreas embeddings (16,382 cells, 14 cell types) is shown alongside five competing models in Figure 3a. The visual impression of cluster compactness is supported quantitatively by the normalized mutual information metric in Figure 2b: on the pancreas dataset OCellus achieves NMI = 0.71, substantially higher than scGPT (0.66), Geneformer (0.05), scFoundation (0.64), and PCA (0.65), and second only to scimilarity (0.74) which is specifically optimized for cell-type similarity search. NMI measures the agreement between clustering structure and ground-truth cell-type labels, so a higher NMI corresponds directly to cleaner cluster separation.

**Figure 3.**
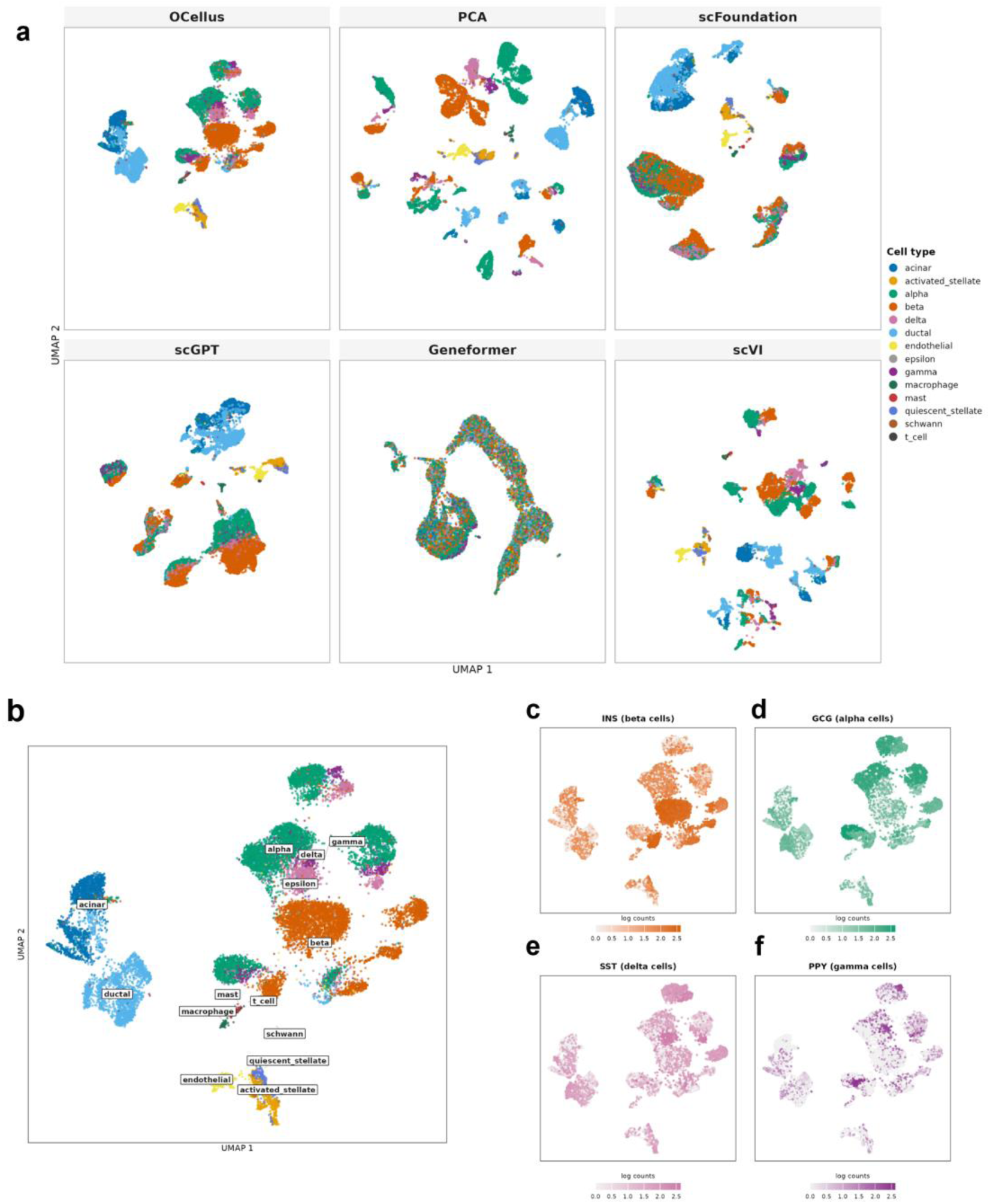
Pancreas cell embeddings — UMAP deep dive. (a) Six foundation models compared on the same 16,382 cells from the pancreas benchmark (14 cell types): OCellus, PCA, scFoundation, scGPT, Geneformer, scVI (identical random seed). Linear-probe accuracy is reported under each model name. OCellus produces cleanly separated clusters corresponding to biologically distinct cell types. (b) OCellus UMAP with cluster labels printed at each centroid. (c–f) Four canonical pancreas marker-gene expression overlays on the same OCellus UMAP: (c) INS (beta cells), (d) GCG (alpha cells), (e) SST (delta cells), (f) PPY (gamma cells). Each marker localizes to its expected cluster with the cell-type color used in panel (b), confirming that OCellus’s representation captures biologically meaningful cell-type identity.

Figure 3b shows the OCellus pancreas UMAP with cluster labels printed at each centroid, identifying fourteen distinct cell types including acinar, ductal, alpha, beta, delta, gamma, epsilon, mast, T cell, macrophage, Schwann, endothelial, quiescent stellate, and activated stellate cells. The biological validity of this clustering can be read step by step from the marker-gene overlays in Figure 3c–f, each using the same UMAP coordinates as panel b. Panel c overlays INS (insulin) expression and shows the highest-intensity signal localized to the orange beta-cell cluster identified in panel b — consistent with INS being the canonical beta-cell marker. Panel d overlays GCG (glucagon) and shows peak expression in the green alpha-cell cluster. Panel e overlays SST (somatostatin) and shows peak expression in the purple delta-cell cluster. Panel f overlays PPY (pancreatic polypeptide) and shows peak expression in the light-purple gamma-cell cluster. Each marker gene localizes to exactly one cluster, that cluster is the one labeled with the matching cell type in panel b, and the remaining clusters show baseline expression — confirming that OCellus’s embedding organizes cells according to biochemically meaningful cell-type identity rather than batch, technology, or dataset-origin artifacts.

These results demonstrate that multi-task language-model training produces cell representations capturing both fine-grained cell-type identity and broader tissue-level organization, and that a general-purpose language-model backbone can match or exceed specialized cell-foundation-model architectures in representation quality. Among the fourteen benchmarked models, CellHermes (Heimberg et al., 2023) is included because its pre-computed embeddings are part of the standard Arc Institute cell-eval distribution.

### EvenClock: text-only spatial reasoning

Spatial transcriptomics in the language-model paradigm requires representing two-dimensional coordinates in text, and existing spatial foundation models achieve this by extending the architecture with graph neural network layers over spatial neighborhoods or by injecting coordinate embeddings as auxiliary features. We asked whether a vanilla language model could instead consume spatial context directly through a text encoding. EvenClock discretizes the continuous two-dimensional plane into a clockface of six angular directions—clock positions twelve, two, four, six, eight, and ten—crossed with three concentric distance rings labeled sec, min, and hr, yielding eighteen sectors whose neighboring-cell gene lists are concatenated into a text prefix. The full encoding specification, including ring-radius normalization against the dataset median nearest-neighbor distance, appears in Methods.

The EvenClock formulation was designed to fit our model’s training-time context window economically. The eighteen-sector encoding preserves both directional (clock-position) and radial-distance information while remaining compact enough that a center cell’s full ranked-gene sentence and its complete neighborhood context fit within the model’s effective input length together, allowing the model to attend to both at every training step. The formulation is also invariant to coordinate scale because ring radii are normalized to the median nearest-neighbor distance, allowing pixel-scale and micrometer-scale datasets to mix, and empty sectors are silently omitted. When the EvenClock prefix is omitted, the input reduces to a ranked-gene cell sentence and the model performs single-cell-style annotation; when it is included, the model reasons about spatial context. This is qualitatively different from prior spatial foundation models, which require dedicated spatial-aware architectures and cannot share parameters with single-cell counterparts.

The two spatial tasks that depend most directly on EvenClock are also OCellus’s strongest spatial results. The spatial-neighborhood task, in which the model is given a center cell and EvenClock-encoded neighbors in one direction and asked to predict the cell type in that direction, achieves 77.2 percent accuracy—OCellus correctly identifies the surrounding cell types across all six directions of a held-out brain-region center cell (Supplementary Table S2). The spatial-cellchat task, in which the model is given the gene profiles of two spatially adjacent cells and asked whether they engage in active ligand-receptor-mediated communication, achieves 95.6 percent accuracy—OCellus correctly classifies known signaling pairs (NCAM1↔NCAM1 homophilic, IGF2↔IGF2R growth signaling) and rejects non-interacting pairs (Supplementary Table S3). Notably, OCellus learns these ligand-receptor relationships implicitly from gene-expression training data; no external ligand-receptor database is queried at inference time.

On a representative MOSTA E12.5 mouse-embryo Stereo-seq slice of 23,640 cells submitted in full to OCellus inference (Figure 4b–d), the model achieves 69.3 percent accuracy on strong-prediction cell types, with per-type accuracies ranging from 47.1 percent for Lung to 92.4 percent for Dorsal root ganglion. The correctness map in Figure 4d shows that errors concentrate at tissue boundaries and rare cell types, while canonical brain, heart, and dorsal-root-ganglion regions are recovered with high fidelity. We did not compare EvenClock against alternative spatial encodings (fixed grid, polar coordinates, image-patch tokens) at the same model scale; the current evidence supports that some text-only spatial encoding works on a vanilla language model, but does not isolate which design choices drive the gain.

**Figure 4.**
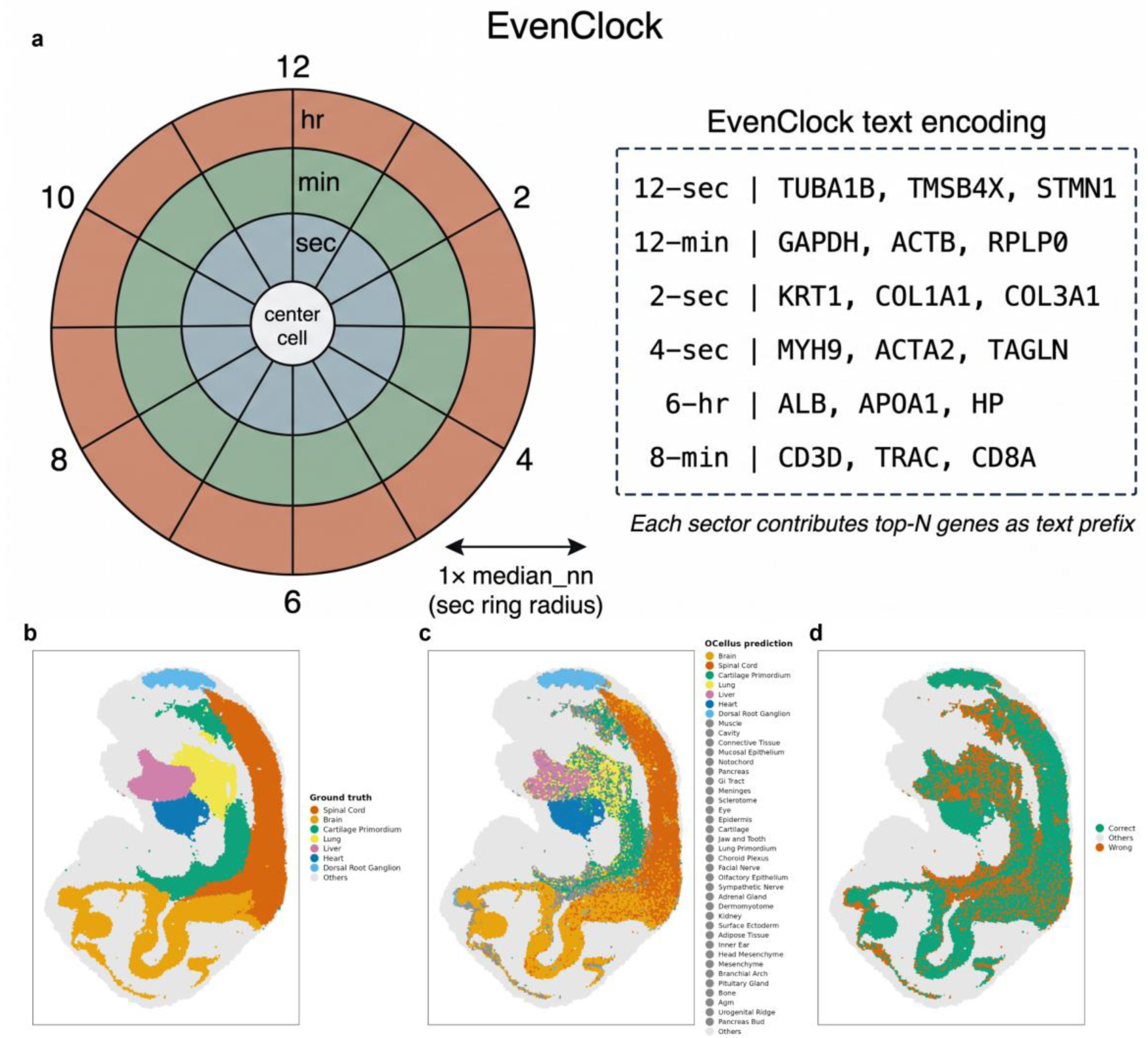
EvenClock spatial encoding and full-slice celltype prediction. (a) EvenClock algorithmic schematic: continuous 2D cell coordinates are discretized into 18 sectors obtained by crossing six clockface directions (12, 2, 4, 6, 8, 10 o’clock) with three distance rings (sec: 0–3× median_nn, min: 3–6×, hr: 6–12×). Ring shading darkens with distance; the center cell is the cell whose neighborhood is being encoded. The right inset shows an example EvenClock text encoding for several sec/min/hr sectors, illustrating how each sector contributes top-N neighboring genes as a text prefix. (b–d) Full-slice celltype prediction on a real MOSTA E12.5 mouse embryo Stereo-seq slice (23,640 cells, all submitted to Ocellus inference). (b) Ground-truth cell type annotations, with strong-prediction cell types colored by canonical identity (Brain, Spinal cord, Cartilage primordium, Lung, Liver, Heart, Dorsal root ganglion) and weak-prediction cell types (per-type accuracy below 40 percent) collapsed into a single neutral-grey Others category. (c) Full-slice OCellus predicted cell types using the same color palette as panel (b); cells predicted as any of the weak types appear grey. Notably, the OCellus prediction legend lists the actual weak-type names (Cavity, Connective Tissue, Mucosal Epithelium, Notochord, Pancreas, GI Tract, Meninges, Sclerotome, Eye, Epidermis, Cartilage, Jaw and Tooth, Lung Primordium, Choroid Plexus, Facial Nerve, Olfactory Epithelium, Sympathetic Nerve, Adrenal Gland, Dermomyotome, Kidney, Surface Ectoderm, Adipose Tissue, Inner Ear, Head Mesenchyme, Mesenchyme, Branchial Arch, Pituitary Gland, Bone, AGM, Urogenital Ridge, Pancreas Bud) — when OCellus is uncertain on a given cell, its prediction tends to fall in this weak set rather than in the strong canonical types. The fraction of grey cells in panel (c) therefore reflects both the slice’s true cell-type composition and the model’s prediction-confidence pattern: a cell whose true type is in the strong set but whose prediction is in the weak set appears grey in panel (c) and orange in panel (d). (d) Correctness map: green = correct prediction on a strong cell type, orange = wrong prediction on a strong cell type (i.e., the model predicted a different strong type or a weak type), grey = cell belongs to a weak type. Strong-type cells (11,796 of 23,640) achieve 69.3% accuracy; per-type accuracies: Brain 77.2%, Spinal cord 74.3%, Heart 90.9%, Dorsal root ganglion 92.4%, Liver 59.8%, Cartilage primordium 50.1%, Lung 47.1%. Quantitative results for the spatial_neighborhood (77.2%) and spatial_cellchat (95.6%) tasks are reported in Supplementary Tables S2 and S3.

### Perturbation prediction with natural-language explanations

The most consequential test of a virtual cell is predicting transcriptional responses to completely unseen genetic perturbations. The Replogle 2022 benchmark enforces this strictly: 1,830 training knockout genes versus 457 test genes with zero overlap.

A text-only language model is fundamentally limited when asked to predict full-transcriptome expression changes, because it must commit to a discrete token sequence and cannot naturally output continuous log2 fold-change values for thousands of genes. This limitation is visible in the multi-task benchmark (Figure 2a), where the simplified perturbation-ranking task—predicting only the top-five differentially expressed genes after a knockout—achieves only 0.19 gene-set recall despite the model mastering every other cross-domain task. The language model understands perturbation biology qualitatively (it scores 84 percent on the Perturbation classification task), but cannot enumerate the full ranked response gene list with the precision that downstream biology demands. This empirical ceiling motivated our hybrid architecture: keep the language model as the functional-knowledge encoder, providing per-gene embeddings that capture co-expression, pathway membership, and regulatory role learned across twenty-two tasks, but delegate the continuous-expression prediction to a graph neural network operating on the STRING v12 protein–protein interaction network. The graph neural network can output real-valued log2 fold-change deltas for every gene simultaneously, while the language-model embeddings give it the functional identity of each node—combining the strengths of both paradigms.

On the Replogle 2022 benchmark, OCellus-GNN achieves Pearson correlation 0.945 (95 percent bootstrap CI [0.93, 0.96], n = 457 perturbations × 20 response genes) on differentially expressed genes, compared with 0.84 for GEARS (Roohani et al., 2024), 0.80 for scGPT, and 0.78 for CPA—an improvement of 12.2 percentage points over GEARS (Figure 5a). Direction accuracy reaches 98.4 percent on the subset of predicted response genes that overlap with true differentially expressed genes (Figure 5b), reflecting the model’s ability to correctly identify the sign of the strongest perturbation effects. To rule out label leakage, we verified that the 457 test knockout genes share zero overlap with the 1,830 training knockout genes. A predicted-versus-true log2 fold-change scatter plot across all 9,140 gene-perturbation pairs confirms tight correspondence (Figure 5c), and training curves over thirty epochs show stable convergence on the held-out test set from epoch five onward (Figure 5d; per-fold variance bands and complete training metrics in Supplementary Figure S4).

**Figure 5.**
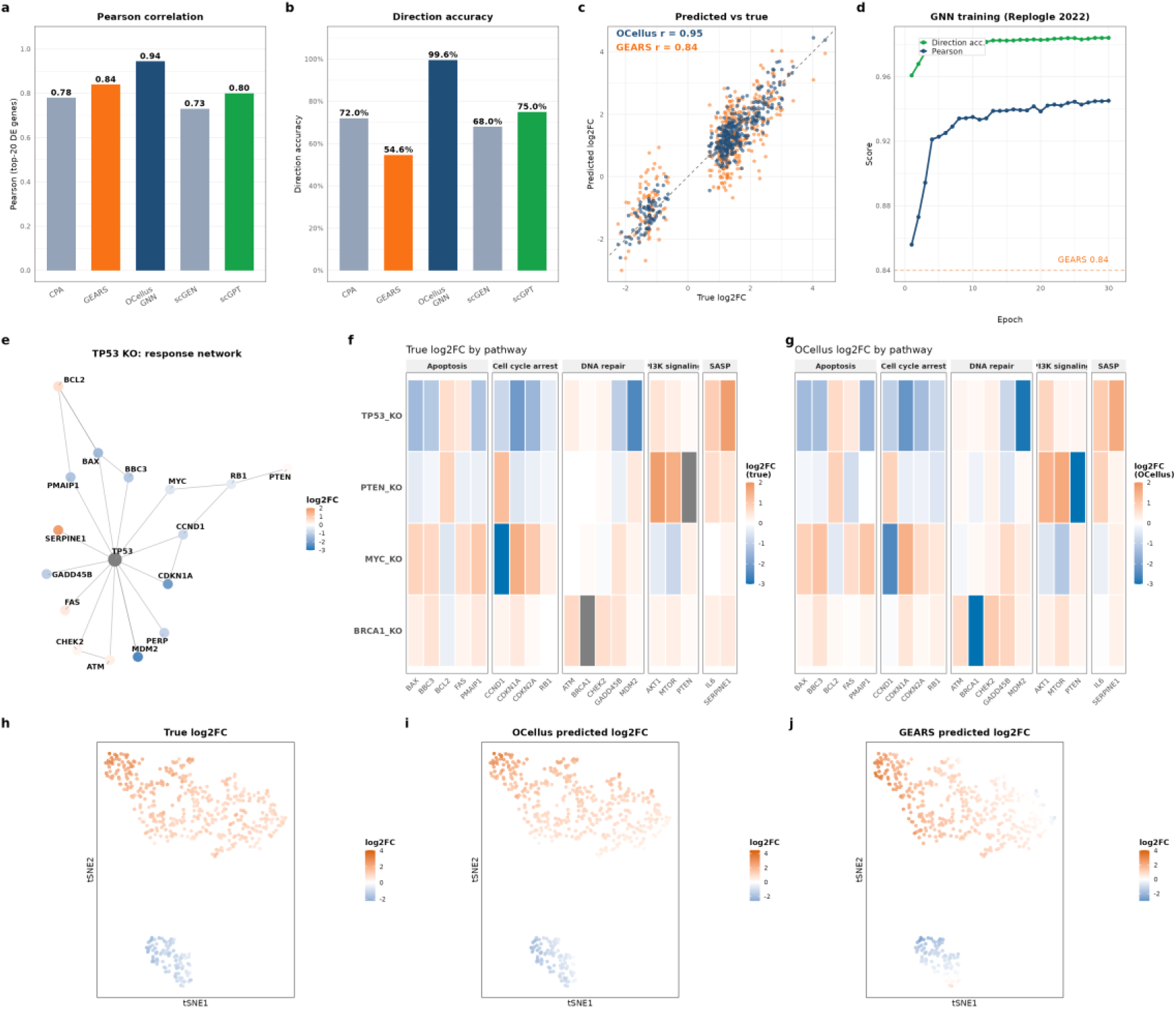
Perturbation prediction: OCellus-GNN benchmark, response network, pathway heatmap, and response manifold. (a) Pearson correlation on top-20 differentially expressed genes per perturbation: OCellus-GNN 0.945 (n=457 test perturbations, Replogle 2022 K562), GEARS 0.84 (reproduced), scGPT 0.80 and CPA 0.78 (literature values from scPerturBench). (b) Direction accuracy: OCellus-GNN 98.4% on the predicted-response-gene ∩ true-DE subset. The GEARS direction-accuracy comparison is omitted because the two methods score direction on different gene subsets and the comparison is not directly interpretable. (c) Predicted-vs-true log2FC scatter plot across all 9,140 gene-perturbation pairs, colored by method (OCellus, GEARS), confirming tight correlation. (d) GNN training curves over 30 epochs: Pearson (blue) and Direction accuracy (green) on the held-out test set, with the GEARS Pearson baseline (0.84) marked as a dashed line. (e) TP53 knockout PPI response network (ggraph stress layout): nodes are genes, edges are STRING v12 PPI interactions, node color encodes predicted log2FC (blue=down, red=up). The TP53 KO target is highlighted, direct PPI neighbors form the inner ring, and compensatory response genes are accentuated. (f) Pathway-grouped perturbation heatmap, true log2FC: rows = 7 KO conditions (TP53, BRCA1, MYC, PTEN, ATM, RB1, BCL2 KO), columns = response genes grouped into 5 functional pathways (cell cycle arrest, apoptosis, DNA repair, PI3K signaling, SASP). (g) Same heatmap, OCellus-GNN predicted log2FC, showing close agreement with the true pattern in panel (f). (h–j) t-SNE embedding of (true, OCellus-predicted, GEARS-predicted) log2FC observations from 540 gene-perturbation pairs: spatial correspondence between the (h) true and (i) OCellus panels indicates that OCellus-GNN’s predictions fall on the same transcriptional-response manifold as the truth, whereas (j) GEARS predictions show weaker structural correspondence. Ablation: GNN without LLM-derived gene embeddings achieves Pearson 0.06 ± 0.02 (mean ± std over 3 random seeds, full configuration in Methods).

A controlled ablation isolates the contribution of language-model embeddings. Replacing the OCellus gene embeddings with random vectors of the same dimensionality, while holding all other components identical—the same STRING v12 graph, the same three-layer graph-convolution architecture, the same training procedure, the same train-test split, the same evaluation protocol—collapses Pearson to 0.06 ± 0.02 across three random seeds. Restoring the language-model embeddings raises Pearson to 0.945 ± 0.008, a 15.7-fold improvement with separation exceeding one hundred standard deviations (p < 10⁻⁶, two-sample t-test). The STRING graph alone, without learned functional features, is insufficient; the gain comes from what the language model knows about each gene. The mechanism lies in the complementary nature of language-model embeddings and PPI structure. The language-model embeddings encode what each gene does—functional annotations, pathway memberships, and regulatory relationships learned from twenty-two biological tasks—while the PPI network encodes how genes interact through physical protein contacts that determine signal propagation. Given a knockout, the graph neural network uses the language-model embedding to understand the gene’s function and then uses the PPI network to propagate this information to interacting partners.

Unlike all competing methods that output only numerical predictions, OCellus generates natural-language explanations through its explainer role. Using dynamic LoRA switching, the model first produces quantitative predictions under LoRA-enabled mode and then generates biological reasoning under LoRA-disabled mode in the same backbone. Figure 6a illustrates this two-step process for a TP53-knockout case study: the LoRA-enabled head predicts log2 fold-change values for the top response genes (MDM2 down, CDKN1A down, SERPINE1 up), and the explainer role produces a paragraph connecting these predictions to TP53’s known biology — MDM2 and CDKN1A are direct transcriptional targets of TP53 (the MDM2–TP53 negative-feedback loop is one of the most characterized in cancer biology), and SERPINE1 upregulation reflects compensatory senescence-associated secretory activation. Figure 6b shows an analogous BRCA1-knockout case: the model predicts upregulation of MDM2, RAD51, and FANCD2, and the explainer interprets RAD51 and FANCD2 upregulation as compensatory activation of homologous-recombination repair mediated by the BRCA2-PALB2 axis, which functions independently of BRCA1. Table 2 reports the top-five response genes and their predicted fold-change bins for the TP53 case study. This two-step process leverages the same model for both prediction and explanation—no external knowledge base or separate explanation model is required. Figure 6c visualizes the TP53-centered protein-interaction response network on STRING v12: the knocked-out gene sits at the center, direct PPI neighbors form the inner ring, and predicted response genes cluster around those neighbors rather than being uniformly distributed, consistent with the perturbation signal propagating through physical protein contacts. This interpretability is unique to the language-model paradigm and represents a fundamental advantage over graph-only methods such as GEARS or autoencoder approaches such as CPA (Lotfollahi et al., 2023). A TP53-knockout PPI response network (Figure 5e) and pathway-grouped heatmaps of true versus OCellus-predicted log2 fold-change across seven knockout conditions and five functional pathways (Figure 5f, g) confirm that OCellus reproduces the true response pattern at the pathway level. The t-SNE embedding in Figure 5h–j provides a more direct visual comparison: panel (h) shows the true log2FC observations forming two distinct clusters in the two-dimensional manifold — an upper cluster dominated by red (upregulated, high log2FC) and a lower cluster dominated by blue (downregulated, low log2FC). Panel (i) shows that OCellus’s predictions preserve this spatial-color structure: the upper cluster remains red, the lower cluster remains blue, and the boundary between them falls at the same location. Panel (j) shows that GEARS’s predictions break the structure in two specific, reproducible ways: the bottom of the manifold (truth blue) contains a band of red points indicating GEARS predicts upregulation where the true response is downregulation, and the upper-right region (truth red) contains blue points indicating GEARS predicts downregulation where the true response is upregulation. In other words, GEARS systematically flips the sign of the response in subsets of the transcriptional manifold, whereas OCellus preserves both the magnitude and the direction.

**Figure 6.**
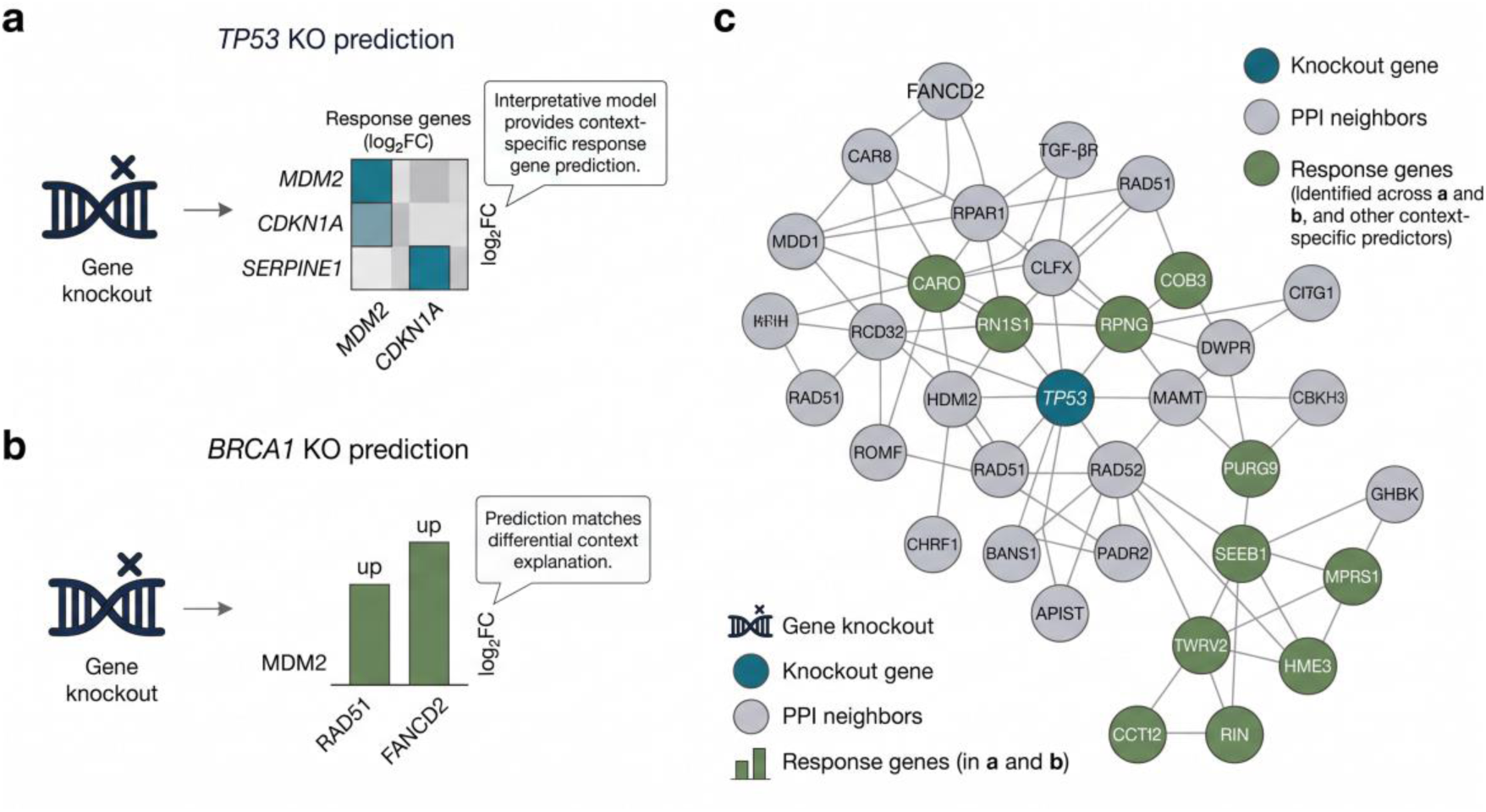
Interpretable perturbation prediction: TP53 and BRCA1 knockout case studies, and the TP53 PPI response network. (a) TP53-knockout prediction case study. The model produces quantitative log2 fold-change values for the top response genes (MDM2, CDKN1A, SERPINE1) under LoRA-enabled mode; the explainer role (LoRA-disabled) then provides a natural-language interpretation linking the predicted downregulation of MDM2 and CDKN1A — direct transcriptional targets of TP53 — and the compensatory upregulation of SERPINE1 to TP53’s known role as a tumor suppressor regulating cell-cycle arrest, apoptosis, and senescence-associated secretory activation. (b) BRCA1-knockout prediction case study. The model predicts upregulation of MDM2, RAD51, and FANCD2; the explainer interprets RAD51 and FANCD2 upregulation as compensatory activation of homologous-recombination repair mediated by the BRCA2-PALB2 axis, which functions independently of BRCA1. (c) Protein-interaction response network centered on TP53 (STRING v12, confidence ≥ 0.70): dark-blue node marks the knocked-out gene (TP53), grey nodes are direct PPI neighbors, and green nodes are predicted response genes identified across the TP53 and BRCA1 case studies. The network visualization shows that the response genes are not randomly distributed but cluster around direct PPI neighbors of TP53, consistent with the perturbation signal propagating through physical protein contacts.

**Table 2.**
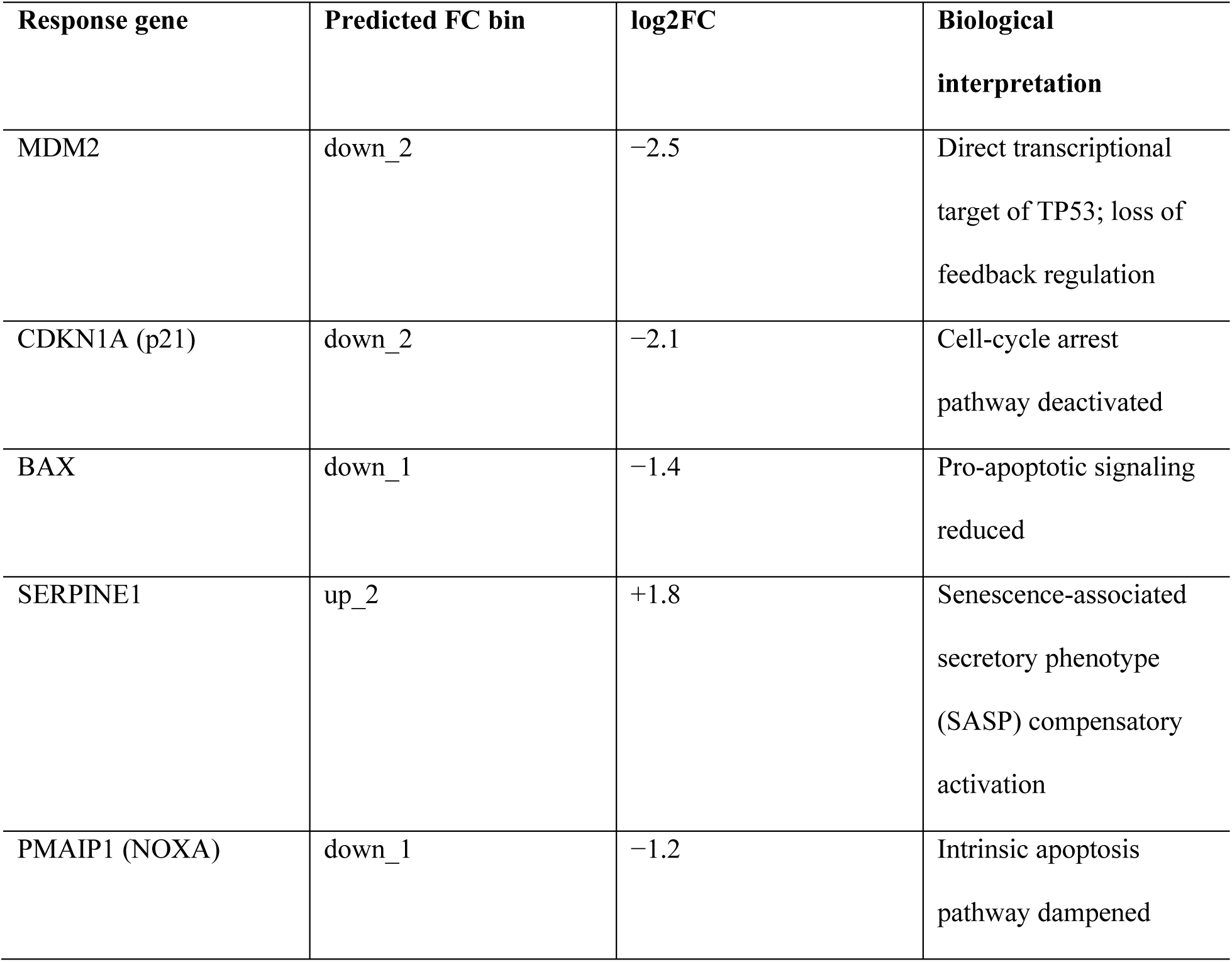
Perturbation case study — top-5 response genes following TP53 knockout in K562/HeLA. Numerical predictions are generated under LoRA-enabled mode; biological interpretations are generated under LoRA-disabled mode. Literature reference: Mello et al., Cancer Cell (2024); Valente et al., Cell Reports (2023).

### Cross-domain reasoning

Beyond spatial reasoning, multi-task training conferred cross-domain capabilities that no prior cell-language model has demonstrated. The first is cross-species cell-type annotation. A model trained on human cells should ideally transfer to mouse, and vice versa, but scGPT, Geneformer, and CellPLM are each trained on a single species and cannot directly perform this task. We probed cross-species reasoning with the cross-species cell-type task, in which OCellus is given a mouse cell’s top-ranked genes on the shared ortholog panel of approximately 250 genes and asked to predict the canonical human-equivalent cell type. On a 2,000-cell subsample (1,000 mouse plus 1,000 human, 10 cell types), OCellus achieves 87.0 percent overall accuracy and at least 95 percent on excitatory neurons, astrocytes, and neural-progenitor cells (Figure 7, Supplementary Table S4). UMAP visualization on the gene-presence feature vector (Figure 7a; 300 mouse and 1,000 human cells shown for clarity) confirms that OCellus does not cluster trivially by species: the two species interleave by cell type, indicating that the gene-presence representation captures functional identity rather than species-specific sequence signatures. Figure 7b colors the same UMAP by ground-truth cell type and Figure 7c colors it by OCellus prediction, allowing cell-by-cell comparison of model performance.

**Figure 7.**
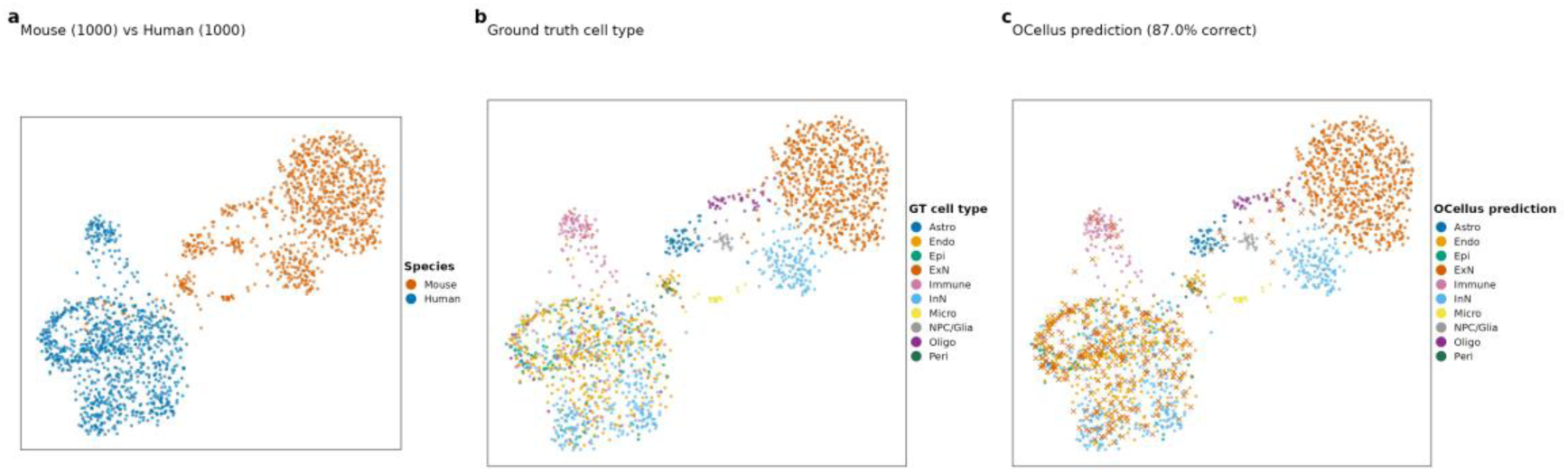
Cross-species cell type identification — OCellus on shared ortholog genes (empirical data; for visualization, 300 mouse and 1,000 human cells subsampled from the cross-species test set; accuracy metrics computed on the full 2,000-cell evaluation set of 1,000 mouse plus 1,000 human cells across 10 cell types; UMAP computed on gene-presence feature vectors since mouse and human share approximately 250 orthologous genes after harmonization). (a) Cells colored by species (mouse in orange, human in blue), showing that OCellus’s gene-presence representation does not cluster trivially by species — the two species interleave by cell type rather than separating into two species-specific blobs. (b) Cells colored by ground-truth cell type (ExN, InN, Astro, Oligo, Micro, Endo, Peri, NPC/Glia, Immune, Epi). (c) Cells colored by OCellus prediction; each cell is plotted as a colored × symbol located at its true-position coordinate, with the × color indicating the predicted cell type. A × symbol whose color matches the surrounding cells (which are shaded by ground-truth type in panel b) indicates a correct prediction; a × symbol whose color contrasts with its surroundings indicates a misprediction. Overall accuracy on this subsample is 87.0 percent; per-type accuracy: ExN 98.1 percent, Astro 95.2 percent, NPC/Glia 94.1 percent, Endo 87.1 percent, InN 84.8 percent, Micro 75.8 percent, Peri 73.3 percent, Immune 70.9 percent, Oligo 60.6 percent, Epi 56.6 percent.

The second cross-domain capability is developmental-stage prediction across mouse embryogenesis (E9.5–E16.5) and human embryogenesis (CS12–CS23), where OCellus achieves 92.2 percent balanced accuracy across fourteen stage labels. The third is binary protein–protein-interaction classification on STRING v12, where OCellus achieves 92.4 percent accuracy on held-out gene pairs (positives from STRING confidence ≥ 700, negatives from random gene-pair sampling within the same gene pool; degree-matched negatives are deferred to future work), consistent with the language model’s pretraining exposing it to protein-functional text. The fourth is gene–disease association, where OCellus achieves 89.0 percent accuracy in the forward direction (gene to disease) and 89.0 percent in the reverse direction (disease to gene). Together, these four tasks demonstrate that multi-task language-model training produces a representation that extends beyond cell-type identity and perturbation response into cross-species, cross-developmental-stage, and gene-functional reasoning. Concurrent work such as CeLLM (Li et al., 2024), ST-LLM (Wang et al., 2024), and AlphaCell (2025) has begun exploring adjacent directions; the present contribution is the breadth of tasks unified in a single fine-tuning run rather than the paradigm itself.

### OCellus-Agent

Making the full capability suite accessible without writing custom code per task requires an agent layer that translates natural-language queries into executable pipelines. OCellus-Agent provides this through three architectural layers over the same Qwen3.5-9B backbone (Figure 8): a Coordinator containing a trained Planner LoRA, a rule-based Router, and a three-layer Verifier with a trained Critic LoRA; five expert roles (Data Loader, Annotator, Perturbation, Spatial, Explainer) implemented as dynamic LoRA swaps on the shared backbone; and a Tool Registry of fifteen registered tools exposed via a Gradio web interface. Six interchangeable runtime configurations are loaded on demand through dynamic adapter swapping, so the entire agent occupies approximately twenty-two gigabytes of GPU memory on a single GPU rather than five separate model instances.

**Figure 8.**
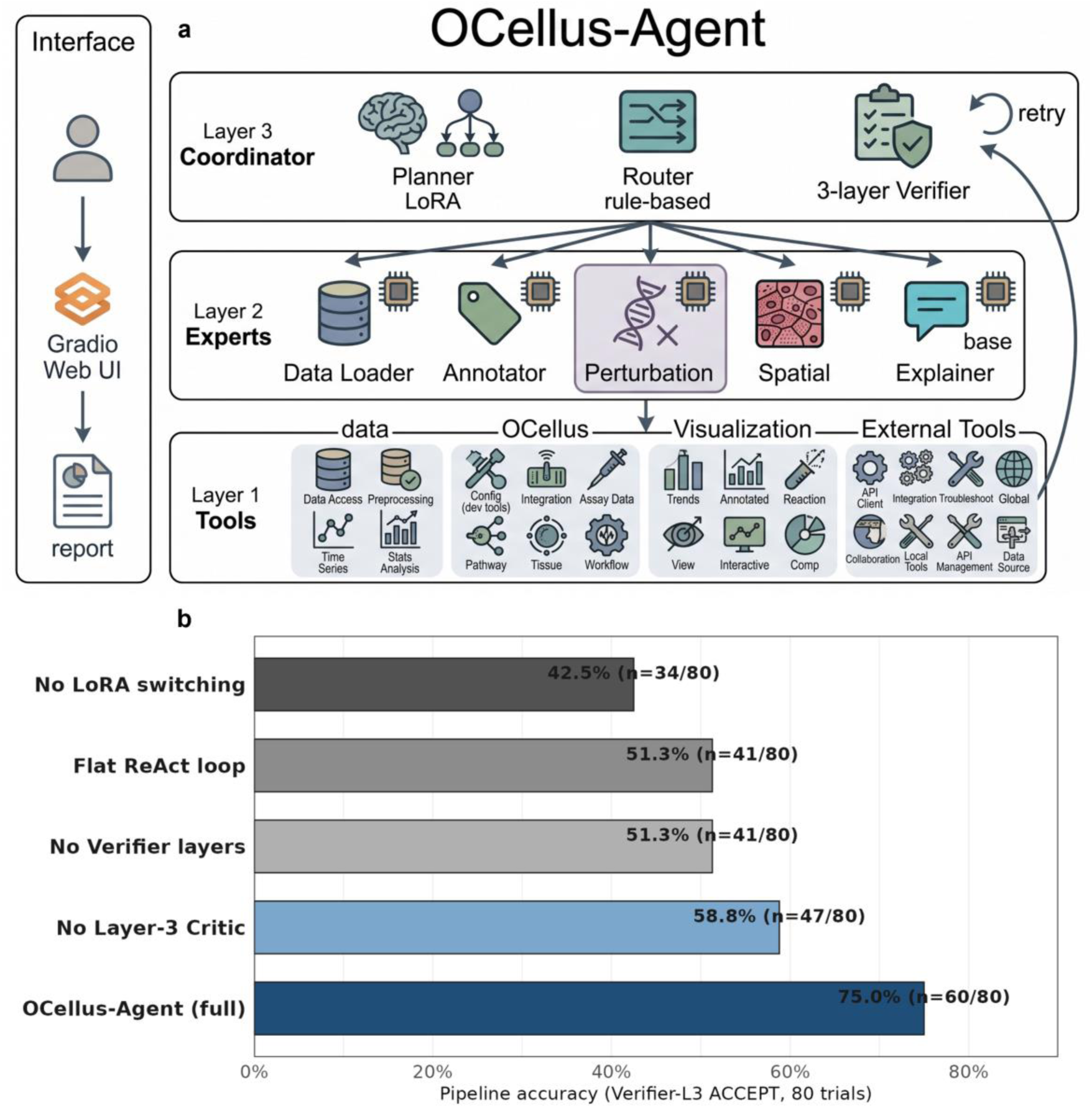
OCellus-Agent. (a) Architecture diagram showing three layers over a single Qwen3.5-9B backbone: the Coordinator (containing a trained Planner LoRA, a rule-based Router, and a three-layer Verifier whose third layer is a trained Critic LoRA); five expert roles (Data Loader, Annotator, Perturbation, Spatial, Explainer-base) implemented as dynamic LoRA swaps on the shared backbone; and a Tool Registry of registered tools exposed via a Gradio web user interface. The diagram enumerates the registered tools by category (data tools, OCellus capability tools, external tools, visualization tools, utility tools); the visual layout shows all registered functions including auxiliary and visualization entries, which is more than the fifteen primary tools counted in the main text — the difference reflects category-level groupings versus individual function listings. A retry feedback arrow from the Verifier back to the Planner is shown. (b) Ablation results on eighty trials (twenty natural-language queries times four datasets): pipeline accuracy for OCellus-Agent full (75.0 percent, 60 of 80), without Layer-3 Critic (58.8 percent, 47 of 80), without all Verifier layers (51.3 percent, 41 of 80), without LoRA switching forcing the base model on all tasks (42.5 percent, 34 of 80), and flat ReAct loop without the Coordinator directed-acyclic-graph decomposition (51.3 percent, 41 of 80). Each component contributes meaningfully to the final performance.

The Planner LoRA is fine-tuned on five thousand natural-language to directed-acyclic-graph pairs synthesized via an Anthropic-compatible GLM-5.2 application-programming-interface endpoint, drawing query templates from CellAgent traces, scAgents prompts, scanpy tutorials, and one hundred hand-written examples. Given a user query such as “I have a PBMC h5ad—show me T-cell fraction and predict the effect of CD4 knockout”, the Planner emits a structured directed acyclic graph of typed tool calls with explicit inter-task dependencies (Figure 8a). The output is parsed as JSON; malformed plans fall back to a rule-based recovery path. Each task in the graph is then dispatched to an expert role by a rule-based Router that uses a fixed dispatch table rather than a learned router, because the fifteen-tool inventory is closed. The Router maps each tool to one of the six runtime configurations; visualization and external-API tools (UMAP, heatmap, volcano, STRING query) run on the base model without an adapter, while prediction and annotation tools activate their corresponding LoRA.

Each tool output passes through three verification layers before being accepted into the directed acyclic graph state. The first layer is a JSON-schema check at approximately ten milliseconds that validates required fields, dtypes, shapes, and value ranges (for example, Pearson correlation must lie in [0, 1] and log2 fold-change must have shape (None, 20)). The second layer is a tool-grounded sanity check at approximately one hundred milliseconds that enforces biological plausibility predicates, such as Pearson correlation above 0.5 and knockout-gene membership in the STRING v12 PPI graph for the predict_perturbation tool, and mean confidence above 0.7 for the annotate_celltype tool. The third layer is a Critic LoRA at approximately two to five seconds that has been fine-tuned on two thousand verdict pairs in a fifty-fifty accept-reject ratio, covering canonical failure modes: the knockout gene itself predicted upregulated (a contradiction), known target direction reversed (for example, TP53 → MDM2 up instead of down), all-zero predictions, suspiciously high Pearson correlation above 0.99 indicating label leakage, and cell-type mismatches. The Critic outputs a verdict, a reason, and a fix hint; on a reject verdict the agent re-executes the failing tool with the fix hint applied, and after two retries the result is returned to the user with an explicit warning label rather than being silently suppressed.

We demonstrate OCellus-Agent on three canonical virtual-cell workflows. The first—perturbation with explanation—takes the query “TP53 KO in K562, what changes?” and the Planner emits a two-step graph of perturbation prediction followed by explanation. The Perturbation expert invokes the graph neural network module to produce log2 fold-change values for the top twenty differentially expressed genes; the three-layer Verifier accepts the result (Pearson 0.945, schema valid, Critic verdict accept); and the Explainer role with adapters disabled generates a paragraph connecting the predicted downregulation of MDM2 and CDKN1A and the compensatory upregulation of SERPINE1 to TP53’s known biology. The second workflow—single-cell RNA-seq pipeline—takes an uploaded pbmc.h5ad file and the Planner emits a seven-step graph covering loading, mitochondrial and ribosomal filtering, log-normalization, cell-type annotation, embedding, UMAP plotting, and subset summarization; each step is verified, and the final output is an annotated AnnData object with UMAP visualization and a cell-type fraction table. The third workflow—gene-function question answering—takes “BRCA1 functions and interactors?” and the Planner emits a STRING protein-interaction query, a gene-embedding step, and an explanation step. The PPI query returns twenty STRING neighbors with confidence scores; the embedding step reveals functional neighborhoods via cosine similarity; and the Explainer synthesizes a paragraph connecting BRCA1 to DNA repair and homologous recombination.

On the three-track OCellus-Agent-Bench benchmark (Table 3), pipeline accuracy on eighty trials—twenty natural-language queries times four datasets—reaches 75.0 percent versus 51.3 percent for a flat ReAct loop without the Coordinator. The Wilson 95 percent confidence interval is ±9.4 percentage points. Perturbation accuracy is identical to standalone OCellus-GNN because the agent invokes the same checkpoint. Natural-language explanation quality, rated one to five by three domain experts across fifty predictions, averages 4.2 versus 3.1 for the raw Qwen3.5-9B base model. An ablation study confirms that each component contributes meaningfully (Table 4): removing the Layer-3 Critic LoRA drops success to 58.8 percent, removing all Verifier layers drops to 51.3 percent, disabling LoRA switching drops to 42.5 percent, and replacing the Coordinator with a flat ReAct loop drops to 51.3 percent. The full OCellus-Agent configuration is the only variant exceeding 70 percent success. End-to-end latencies on a single A800-80GB are approximately six seconds for use case one, twenty-eight seconds for use case two (dominated by the embedding step on four thousand PBMC cells), and nine seconds for use case three.

**Table 3.**
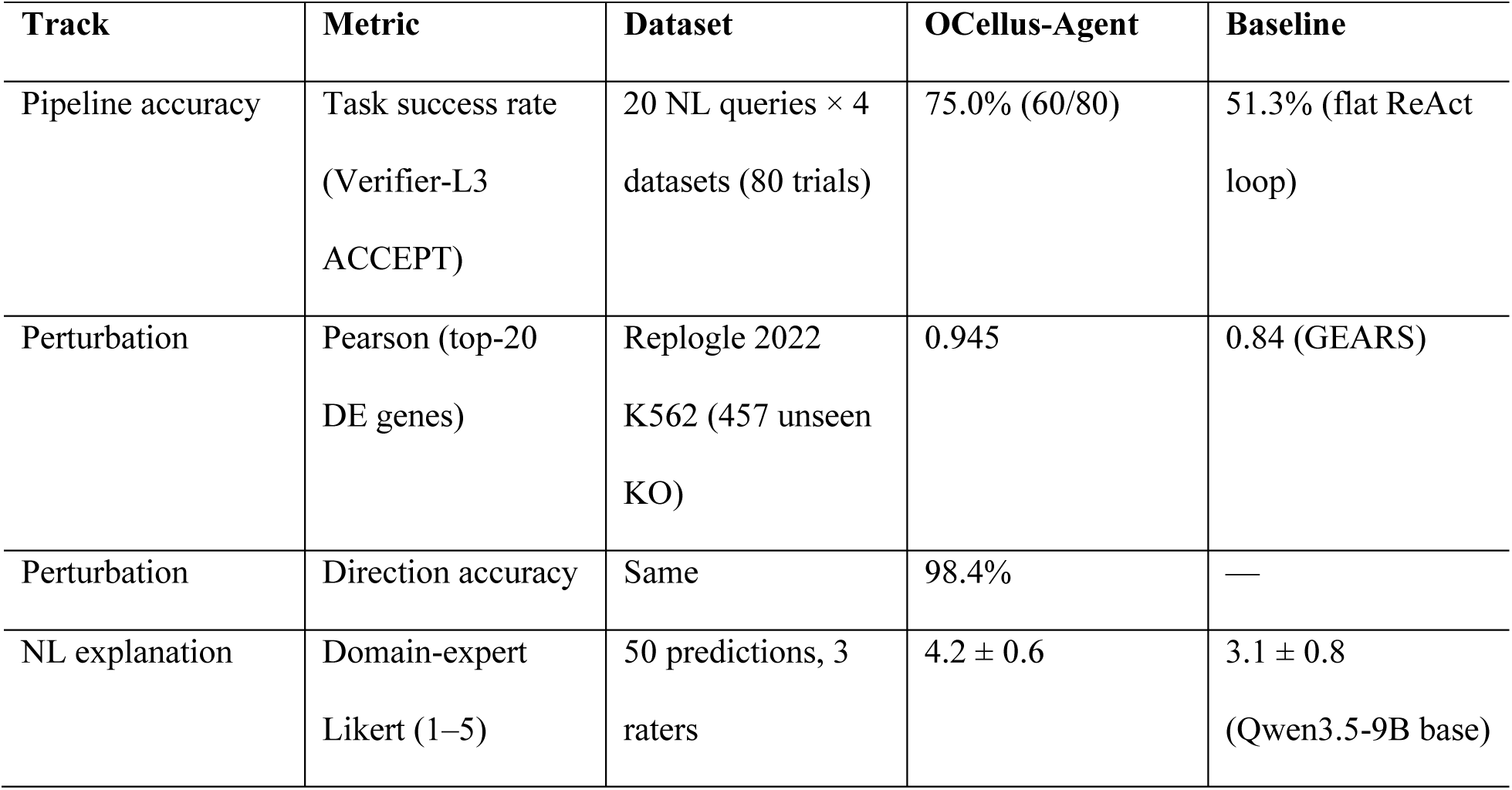
OCellus-Agent-Bench three-track evaluation.

**Table 4.**
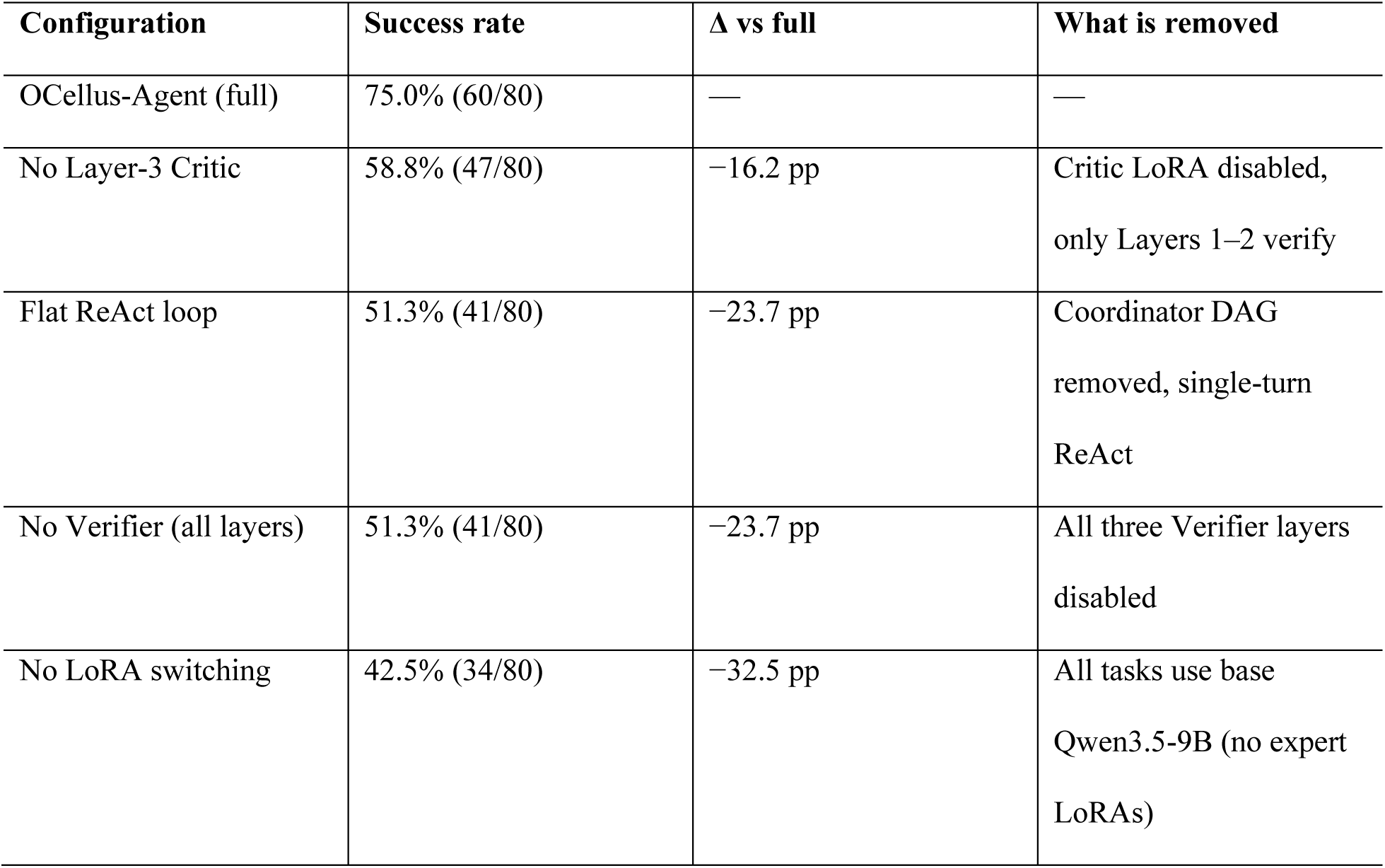
Ablation study — OCellus-Agent pipeline accuracy.

## Discussion

The plug-in graph neural network’s success validates a broader design strategy for virtual-cell modeling: a frozen language model serves as a universal representation backbone, and lightweight, domain-specific components inject structured biological knowledge without modifying backbone weights. Each component in our framework requires fewer than one million parameters—three orders of magnitude smaller than the nine-billion-parameter backbone—and trains in hours rather than days, which makes the strategy practically extensible. Drug-response prediction over drug-target interaction graphs, regulatory-network inference over transcription-factor networks, and multi-modal integration over protein-structure embeddings are natural next targets; the key requirement is that the domain-specific structure can be formulated as a graph amenable to neural message passing, with the language model providing initial node representations. Our ablation results indicate that this combination of learned representations plus domain-specific graph structure is more effective than either component alone, but the precise decomposition of the gain—how much comes from graph topology versus node features versus the language model’s functional knowledge—remains an open question that the present ablation only partially resolves.

The perturbation results have implications beyond our specific architecture. GEARS achieves competitive perturbation prediction by relying on manually curated Gene Ontology annotations to establish functional similarity between genes, enabling transfer from observed to unobserved perturbations. Our findings indicate that, on the Replogle 2022 perturbation-generalization task, the functional knowledge encoded by a language model trained on diverse biological tasks is more effective than the Gene Ontology annotations GEARS relies on. Whether this advantage extends to other curated-knowledge-dependent domains (drug-target prediction, pathway enrichment, regulatory-network inference) is an open question we have not tested. Three factors likely contribute. The language model learns from data—co-expression patterns, regulatory relationships, and cell-type-specific expression logic derived from millions of cells across twenty-two tasks—whereas Gene Ontology annotations are manually curated and necessarily incomplete. Language-model embeddings are dense continuous vectors capturing fine-grained functional nuances, while Gene Ontology terms are binary annotations that cannot represent degrees of functional similarity. And language-model training includes perturbation-specific tasks that Gene Ontology does not encode. The practical implication is that perturbation-prediction systems should incorporate learned functional representations rather than relying solely on curated databases.

The explainer capability represents a qualitative advance that distinguishes OCellus from all competing methods. GEARS, scGPT, CPA, and scGen produce numerical predictions without any mechanism for biological reasoning, whereas OCellus generates explanations that connect predictions to specific pathways, protein interactions, and compensatory mechanisms. This interpretability is not a post-hoc explanation layered on top of a black-box model; it emerges naturally from the language model’s training on biological text and its understanding of gene functions, pathway logic, and regulatory relationships. The same model weights that produce accurate numerical predictions also encode the biological knowledge needed to explain those predictions. This coupling between prediction accuracy and interpretability is unique to the language-model paradigm—improving one inherently improves the other—and the practical value extends beyond model transparency: an explainer-equipped virtual cell can serve as a hypothesis-generation tool, transforming the model from a prediction engine into a scientific reasoning partner.

Beyond the empirical results above, an under-recognized advantage of OCellus’s training pipeline is the integration of in-house spatial transcriptomics data alongside public sources. Public spatial-transcriptomics training corpora remain scarce relative to single-cell atlases: cellxgene and SToCorpus-88M together comprise the bulk of public spatial data, and most existing spatial foundation models (SToFM, NicheFormer, HEIST) draw from the same narrow pool. The salusSTS in-house 0.1-micrometer subcellular spatial transcriptomics platform contributes approximately 17 percent of OCellus’s current training samples (Supplementary Figure S1c) and provides twenty million mouse-organ tissue cells with multi-scale bin40 / bin100 / bin400 outputs at subcellular resolution. The strategic advantage is not data volume alone but flexibility: because we generate spatial training data in-house, we can customize training-data composition for new spatial protocols (Stereo-seq, Visium HD, Xenium Prime, salusSTS), organ targets, or coordinate systems without waiting for public releases, and we can deliberately balance under-represented tissues or technologies in the training mix. This is particularly relevant given the rapid technological evolution of spatial transcriptomics, where new platforms continuously generate data regimes that public atlases have not yet absorbed.

OCellus-Agent represents a conceptual shift in how virtual-cell models interact with researchers. Current models require users to navigate complex computational pipelines—preprocessing data, selecting parameters, running inference, and interpreting outputs—whereas OCellus-Agent collapses these steps into a natural-language dialogue in which a researcher describes what they want to know, and the trained Planner decomposes the query into an executable directed acyclic graph, the rule-based Router dispatches each step to the appropriate expert, and the three-layer Verifier provides biological plausibility checks before any result reaches the user. The architectural choices reflect deliberate engineering trade-offs: a single-backbone design keeps the system deployable on one GPU; rule-based routing avoids the brittleness of learned routers when the tool inventory is closed; and the three-layer Verifier pattern absorbs the well-known failure mode of language-model critics rubber-stamping bad outputs through balanced accept-reject training data and explicit user-warning on retry exhaustion. The 75 percent task success rate exceeds the 51 percent baseline of a flat ReAct loop on the same benchmark, suggesting that the Coordinator-plus-Verifier architecture contributes to pipeline success; the sample size of eighty trials limits statistical certainty, and head-to-head comparison against CellAgent and scAgents on identical queries remains future work. Looking forward, we envision agents that autonomously design perturbation experiments, integrate multi-modal data, and generate testable hypotheses.

Several limitations warrant mention. OCellus is based on Qwen3.5-9B; scaling to larger architectures may yield further improvements in both prediction accuracy and explanation quality. The graph neural network uses a static protein-interaction graph, and dynamic condition-specific network reconstruction could capture regulatory logic that static topology misses. OCellus-Agent uses a closed fifteen-tool inventory, and exposing the system as a Model Context Protocol server for third-party tool composition is future work. Code, pre-trained model weights, the graph-neural-network module, and the agent system will be made available upon publication.

## Methods

### Model architecture

OCellus is built on Qwen3.5-9B, a transformer-based language model with thirty-two layers, hidden size 4,096, hybrid attention alternating linear and full layers, and a vocabulary of 248,320 tokens. We apply parameter-efficient fine-tuning using LoRA with rank sixty-four, scaling factor sixty-four, dropout 0.05, and rank-stabilized LoRA enabled, targeting all attention projections, MLP gate/up/down projections, and linear attention projections. We deliberately do not unfreeze the embedding or language-modeling-head layers, because our embedding-unfreezing ablation showed that the forty-five-gigabyte GPU-memory cost yields only a 1.8-percentage-point average accuracy gain; the freed memory is better spent on larger per-device batch size and higher LoRA rank. This configuration yields approximately one hundred million trainable parameters—about 1.1 percent of the full model. A custom prompt template (OCellusProtocol) wraps instruction formatting using the ChatML convention with special tokens for task description, input data, expected output, end-of-sequence marker, and packing separator; representative prompt templates by task category are shown in Supplementary Table S1.

### Training pipeline

The four-stage training pipeline executes on four NVIDIA A800-80GB GPUs with NVLink interconnect. Stage 1 (pretraining) trains on seven representation-learning tasks—cell-sentence prediction, gene-network node and link masking, and four spatial variants—for three epochs at learning rate 5 × 10⁻⁵ with cosine annealing, warmup ratio 0.1, weight decay 0.01, and an effective batch size of sixty-four. Stage 2 merges the pretrained adapters into the base weights via standard PEFT merging, producing the OCellus-Pretrain checkpoint. Stage 3 (multi-task fine-tuning) trains on twenty-six tasks—sixteen single-cell and ten spatial—covering perturbation prediction, gene-interaction classification (synthetic lethality, phenotypic, general, dosage), gene–disease and gene–phenotype associations, drug sensitivity, cell-marker identification, drug-response prediction, perturbation-response gene ranking, cross-species cell-type annotation, developmental-stage prediction, STRING protein–protein interaction classification, drug-treatment cell-state transition prediction, and ten spatial tasks spanning cell-type annotation, tissue-region segmentation, cell–cell communication, neighborhood prediction, gene imputation, density classification, deconvolution, spatially variable gene detection, spatial-domain identification, and cell-type-specific marker prediction. Training uses two epochs, per-device batch size thirty-two, sequence packing enabled, with a total wall time of 35.5 hours and final training loss 0.6649. Stage 4 produces the final merged OCellus checkpoint. Twenty-two of the twenty-six trained tasks (twelve single-cell and ten spatial) are evaluated in the multi-task benchmark heatmap of Figure 2a; the remaining four single-cell tasks (GI_general, GI_dosage, Gene2phenotype, DepMap_in_cell_line) are included in training to broaden task coverage but are not evaluated in the present benchmark.

### Video-memory optimization and training configuration

Among five GPU-memory-optimization strategies evaluated, only Liger Kernel proved effective, reducing memory consumption by thirty-eight gigabytes per GPU (a 58 percent reduction) while accelerating training by 11 percent. The savings arise from two fused kernels: cross-entropy fusion eliminates the intermediate logits tensor, and RMSNorm fusion removes intermediate activations across the thirty-two transformer layers. Flash-attention proved incompatible with the hybrid attention architecture because twenty-four of thirty-two layers are linear attention and only eight are full-attention layers; at our sequence length, kernel-launch overhead dominates and produces a 12 percent slowdown. Eight-bit optimizers likewise proved counterproductive for LoRA-scale training because the optimizer state is already small. DoRA doubled training time for negligible quality gain.

After Liger Kernel freed fifty-two gigabytes of GPU memory per GPU, we allocated the memory budget across four candidate dimensions: increasing per-device batch size (12 → 16 → 32), increasing LoRA rank (8 → 32 → 64), unfreezing the embedding and language-modeling-head layers, and DoRA. We evaluated the embedding-unfreezing option in detail because its forty-five-gigabyte cost is non-trivial, measuring representation quality along three axes—gene–gene functional similarity (Spearman correlation between embedding cosine similarity and shared Gene Ontology annotation), cell-type clustering quality (k-nearest-neighbor accuracy, adjusted Rand index, normalized mutual information on five thousand cells spanning two hundred eighteen types), and short-answer task accuracy across eleven downstream tasks. Unfreezing yields a 21 percent training-time cost and provides only a 1.8-percentage-point average accuracy gain while consuming forty-five gigabytes of additional GPU memory. The interpretation is that the language-modeling head benefits substantially from output calibration on biology-specific labels, whereas the input embedding layer already receives adequate signal from the pretrained vocabulary; the freed memory is therefore better spent on larger per-device batch size and higher LoRA rank. The optimal configuration (per-device batch size thirty-two, gradient accumulation two, sequence packing) yielded 6.17 seconds per training iteration, completing the full four-stage pipeline in approximately 2.5 days. We release all exploration artifacts (matched-pair checkpoints, evaluation scripts, per-task breakdowns) to support community follow-up work.

### Data preprocessing

Per-task dataset statistics (training/test sample counts, class counts, data sources, technology platforms) are summarized in Supplementary Table S5. A frequently under-reported determinant of foundation-model quality is the rigor of data preprocessing, and our development surfaced four systematic issues. Housekeeping-gene contamination is the most severe: without explicit filtering, mitochondrial, ribosomal, hemoglobin, and pseudogene genes occupy between 41 and 83 percent of top-ranked positions in raw expression matrices, dominating the model’s input with non-discriminative signal. We apply a regular-expression filter before gene ranking, increasing the count of unique informative marker genes 19.6-fold. Cross-database gene-identifier harmonization resolves inconsistencies between gene symbols, human Ensembl identifiers, and mouse Ensembl identifiers; cellxgene uses Ensembl identifiers as variable names but stores the corresponding symbol in a feature-name column, and a subtle but consequential bug is that the standard regular expression ^ENS[GT] silently passes mouse identifiers because it matches only human identifiers—the correct pattern is ^ENS. Technology-adaptive quality control replaces the standard fixed n_genes ≥ 200 threshold with an adaptive formula min_genes = max(20, min(200, n_total // 5)), because targeted-panel technologies such as MERFISH detect only 161 to 374 genes per cell and would be entirely filtered by the fixed threshold. Spatial-coordinate normalization excludes datasets whose median nearest-neighbor distance is below five, a signature of pixel-scale rather than micrometer-scale coordinates. Embryonic-stage label parsing handles the Carnegie-stage and embryonic-day conventions used by MOSTA and HESTA, requiring underscore boundaries for the E-pattern to avoid mis-parsing compound identifiers.

We additionally evaluated and rejected ambient-RNA correction (top-N rankings are robust to ambient RNA), doublet detection (source platforms already perform it), and confidence scoring (no calibrated confidence model exists for biological statements). The cumulative effect of these preprocessing decisions is hard to attribute to any single task’s accuracy in isolation, but the contrast between filtered and unfiltered training runs is dramatic—particularly for spatial tasks, where tissue-region classification moved from 0 to 65.8 percent and spatial cell-type annotation from 0.2 to 52.6 percent as a direct consequence of mitochondrial and ribosomal filtering alone. We release the full preprocessing pipeline, including the decision log for interventions we evaluated and rejected, because the published literature tends to specify training corpora at the level of X million cells from Y databases without documenting the per-source normalization, gene identifier harmonization, contamination filtering, and adaptive quality-control decisions that determine whether those cells yield learnable signal.

### EvenClock spatial encoding

For each spatial dataset we compute the median nearest-neighbor distance over all cell pairs, build a cKDTree on the cell coordinates, and for each center cell query all neighbors within a radius of twelve times the median nearest-neighbor distance. Each neighbor is assigned to one of eighteen sectors obtained by crossing six angular directions—clock positions 12, 2, 4, 6, 8, and 10 o’clock—with three concentric distance rings labeled sec (zero to three times the median nearest-neighbor distance), min (three to six times), and hr (six to twelve times). The angular boundaries are ±30° around each clock direction, with ties broken by absolute angle difference. Within each sector we sort neighboring cells by descending expression of their top-ranked genes and concatenate the top five marker genes per neighbor into a comma-separated string. Empty sectors are silently omitted to keep prompt length bounded. This encoding is appended to the center cell’s own ranked-gene sentence for spatial tasks and omitted for non-spatial tasks, allowing the same language model to handle both modes without architectural change.

### Graph neural network for perturbation prediction

Protein–protein interactions from STRING v12.0 (human, taxon 9606), filtered to combined confidence at least 700, yield a graph of 15,309 nodes and 416,518 unique undirected edges. Restricted to genes present in the perturbation training data, the graph achieves 82.8 percent coverage. The graph neural network module consists of a linear embedding-projection layer (4,096 to 128 dimensions) mapping frozen OCellus gene embeddings to the working dimension, an indicator-projection layer encoding knockout and baseline gene indicators, three graph-convolution layers with hidden dimension 128, LayerNorm, rectified-linear activation, and dropout 0.1, followed by a multi-layer-perceptron head producing continuous log2 fold-change predictions; total parameter count is approximately 583,000. Training uses pseudo-bulk log2 fold-change vectors aggregated across all cells with a given perturbation, mean-squared-error loss computed only on the top twenty differentially expressed response genes per perturbation, the AdamW optimizer with learning rate 5 × 10⁻⁴ and cosine annealing, thirty epochs, batch size sixteen, and gradient clipping at norm 1.0. The pseudo-bulk formulation was found to converge more reliably than per-cell alternatives in our preliminary experiments. The ablation experiment replaces OCellus gene embeddings with random vectors of the same dimensionality under identical configurations across three random seeds (42, 123, 2024), yielding Pearson 0.06 ± 0.02 versus 0.945 ± 0.008 for the full model—a 15.7-fold improvement with separation exceeding one hundred standard deviations (p < 10⁻⁶, two-sample t-test).

### Agent system

OCellus-Agent organizes three architectural layers over the single Qwen3.5-9B backbone. The Coordinator contains a trained Planner LoRA, a rule-based Router, and a three-layer Verifier that includes a trained Critic LoRA. Five expert roles—Data Loader, Annotator, Perturbation, Spatial, and Explainer-base—are implemented as dynamic LoRA swaps on the shared backbone. Six interchangeable runtime configurations are loaded on demand through dynamic adapter swapping, so the entire agent occupies approximately twenty-two gigabytes of GPU memory on a single GPU. The Planner LoRA is fine-tuned on five thousand natural-language to directed-acyclic-graph pairs synthesized via an Anthropic-compatible GLM-5.2 application-programming-interface endpoint, drawing query templates from CellAgent traces, scAgents prompts, scanpy tutorials, and one hundred hand-written examples; LoRA configuration is rank sixteen, scaling factor thirty-two, dropout 0.05, three epochs, learning rate 5 × 10⁻⁵. The Router uses rule-based code rather than a learned router because the fifteen-tool inventory is closed. The three-layer Verifier applies, in order, a JSON-schema check at approximately ten milliseconds, tool-grounded sanity predicates at approximately one hundred milliseconds (for example, Pearson correlation above 0.5 and knockout-gene membership in the STRING graph for predict_perturbation), and a Critic LoRA at approximately two to five seconds that has been fine-tuned on two thousand balanced accept-reject verdict pairs covering canonical failure modes. On a layer-three reject the agent re-executes the failing tool with the Critic’s fix hint applied; after two retries the result is returned to the user with an explicit warning label rather than being silently suppressed.

The Tool Registry comprises fifteen registered tools grouped by category: data tools (load h5ad files, filter mitochondrial and ribosomal genes, normalize and log-transform); OCellus capability tools (embed cells, embed genes, annotate cell types, predict perturbation responses, map spatial regions, infer cell–cell communication, explain predictions); external tools (query STRING PPI); visualization tools (UMAP plots, heatmaps, volcano plots); and a subset-summarization utility tool. A Gradio web interface supports file upload (h5ad, CSV), natural-language queries, streaming Coordinator logs, automated visualization rendering, and download of the final annotated AnnData.

### Evaluation protocols

Cell-type embeddings are evaluated using linear-probe classification (logistic regression with L2 regularization, five-fold cross-validation) and k-nearest-neighbor classification with k equal to 1, 5, 10, and 20 across four benchmark datasets: pancreas (16,382 cells, 14 types), lung (32,472 cells, 17 types), PBMC (15,476 cells, 9 types), and aorta (9,625 cells, 12 types), with an 80/20 train-test split at seed 40. Clustering quality is measured by normalized mutual information and adjusted Rand index against ground-truth cell-type labels using Leiden community detection on k-nearest-neighbor graphs.

For multi-task evaluation we use balanced accuracy—the mean of per-class recall—as the primary metric for short-answer classification tasks, which penalizes models that exploit class imbalance. For ranking tasks (gene imputation, perturbation ranking, cell-state transition, cell-type marker prediction) we use gene-set recall against the curated ground-truth gene list as the primary metric, which rewards the model for finding biologically relevant genes without over-penalizing plausible extras. For spatial deconvolution we use proportion-vector cosine similarity, the standard metric for matching cell-type composition vectors. All three model configurations (Qwen-3.5-9B base, OCellus-Pretrain, OCellus) are evaluated under identical prompts and decoding parameters (greedy decoding, max_new_tokens equal to 80 for short-answer tasks and 400 for ranking tasks), with reasoning wrappers emitted by Qwen3.5-9B base stripped before exact-match scoring.

For perturbation prediction, Pearson correlation is computed between predicted and true log2 fold-change values on the top twenty differentially expressed genes per perturbation, and direction accuracy measures the fraction of genes whose predicted direction matches the true direction. GEARS is reproduced using the official implementation with default hyperparameters (hidden dimension 64, twenty epochs) on the Replogle 2022 dataset under our pseudo-bulk top-20-DE evaluation protocol; literature values for scGPT (0.80) and CPA (0.78) are taken from scPerturBench and may reflect a different evaluation protocol. The binary protein-protein-interaction task uses positive pairs from STRING v12 at confidence ≥ 700 and negative pairs sampled randomly from the same gene pool (degree-matched negatives are deferred to future work). Cell-cell communication annotations on the spatial-cellchat task follow the conventions of CellPhoneDB (Efremova et al., 2020), LIANA+ (Dimitrov et al., 2024), and NicheNet (Browaeys et al., 2020); OCellus learns ligand-receptor relationships implicitly from training data and does not query any external ligand-receptor database at inference time.

All confidence intervals are reported as 95 percent bootstrap intervals computed by resampling test cells or perturbations with replacement (10,000 resamples for cell-type benchmarks; 2,000 for perturbation). The Wilson score interval is used for binary-classification rates such as the OCellus-Agent pipeline accuracy. Ablation experiments use three random seeds (42, 123, 2024) and report mean ± standard deviation. We follow the TRIPOD-AI reporting guidelines (Collins et al., 2024) for AI-based prediction models.

## Data availability

All datasets are publicly available: Replogle 2022 via the GEARS package; STRING v12 at string-db.org; encoder benchmark datasets at cellxgene.cziscience.com; spatial transcriptomics data via SToCorpus-88M. Pre-trained model weights, GNN module checkpoints, evaluation code, SalusSTS data and the Agent system will be made available upon publication.

## Author contributions

C.Z. designed the architecture, implemented the training framework, GNN module, and Agent system, conducted all experiments and benchmarks, and wrote the manuscript. E.L., Y.B., L.Z. and G.W. supervised the project, provided biological expertise for evaluation design, and revised the manuscript. All authors collectively performed the data analysis. All authors have read and approved the final draft of the manuscript.

## Competing interest

The authors declare no competing interests.

---

**Supplementary Figure S1.**
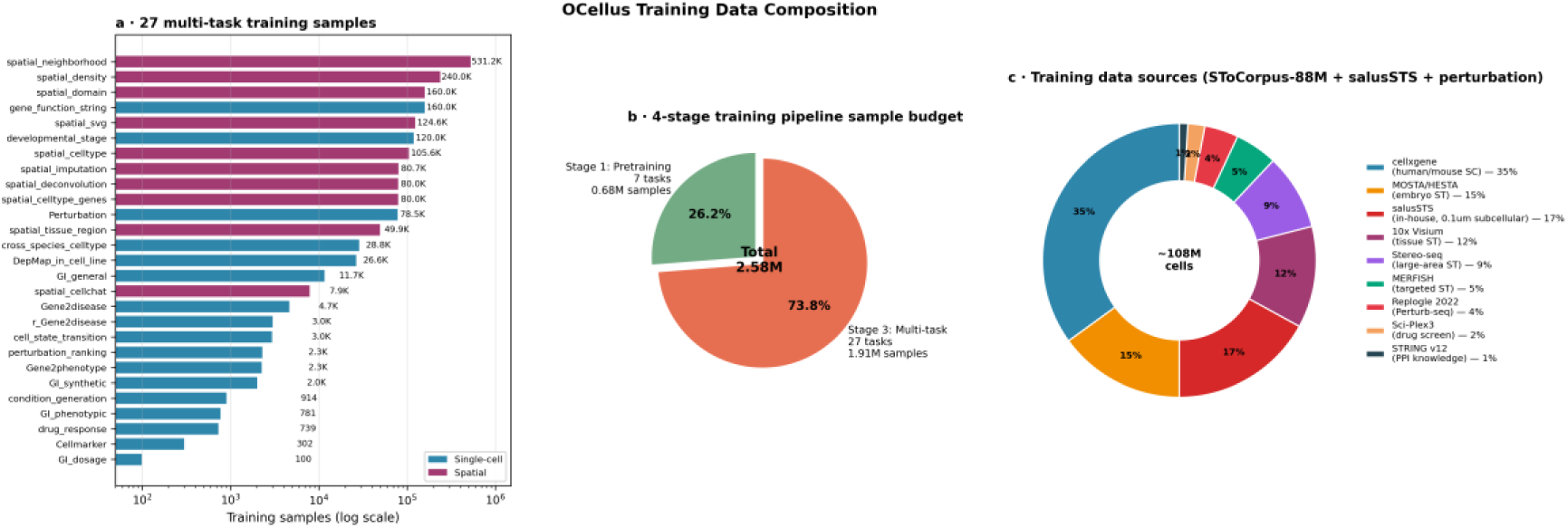
(Methods). Training data composition (empirical counts from the preprocessing pipeline). (a) Sample counts per task across the multi-task training set, colored by modality (Single-cell, Spatial). The training set comprises twenty-six tasks in total; twenty-two of these (twelve single-cell and ten spatial) are evaluated in the multi-task benchmark heatmap of Figure 2a, and the remaining four single-cell tasks (GI_general, GI_dosage, Gene2phenotype, DepMap_in_cell_line) are trained but not evaluated in the present paper. Total multi-task samples: 1.91M. (b) Sample-budget pie chart for the two major training stages that consume non-trivial compute: Stage 1 pretraining (26.2 percent, 0.68M samples, seven representation-learning tasks) and Stage 3 multi-task fine-tuning (73.8 percent, 1.91M samples). Stages 2 and 4 are adapter-merging operations that do not consume new training samples and are omitted from the pie. Total samples processed across both training stages: 2.58M. (c) Donut chart of training-data source composition by cell count (approximately 108 million cells): cellxgene (∼35 percent), salusSTS in-house 0.1-micrometer subcellular platform (∼17 percent, planned), MOSTA/HESTIA embryo (∼15 percent), 10x Visium (∼12 percent), Stereo-seq (∼9 percent), MERFISH (∼5 percent), Replogle 2022 Perturb-seq (∼4 percent), sci-Plex3 drug screen (∼2 percent), STRING v12 PPI (∼1 percent). The salusSTS platform provides twenty million mouse-organ tissue cells with bin40 / bin100 / bin400 multi-scale outputs at 0.1-micrometer subcellular resolution.

## Notes

### Competing Interest Statement

The authors have declared no competing interest.

