## Supplementary Figures for "OCellus: A Language-Model Framework for Single-Cell, Spatial, and Perturbation Biology with Natural-Language Reasoning"

#### Supplementary Figure S1

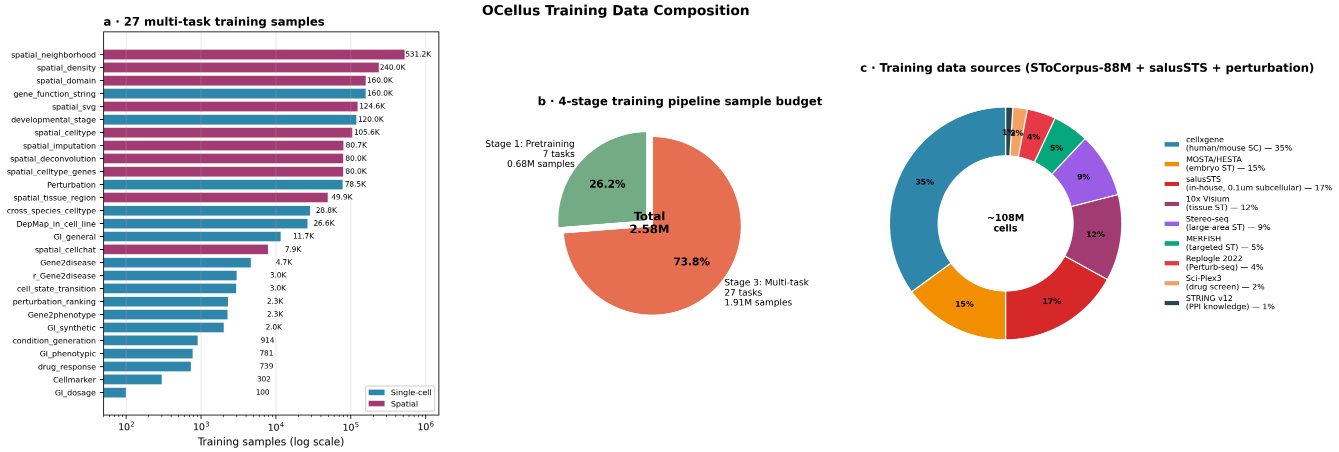


**Supplementary Figure S1. Training data composition** (empirical counts from the preprocessing pipeline). (a) Sample counts per task across the multi-task training set, colored by modality (Single-cell, Spatial). The training set comprises twenty-seven tasks in total; twenty-two of these (twelve single-cell and ten spatial) are evaluated in the multi-task benchmark heatmap of Figure 2a, and the remaining five single-cell tasks (condition_generation, GI_general, GI_dosage, Gene2phenotype, DepMap_in_cell_line) are trained but not evaluated in the present paper. Total multi-task samples: 1.91M. (b) Sample-budget pie chart for the two major training stages that consume non-trivial compute: Stage 1 pretraining (26.2 percent, 0.68M samples, seven representation-learning tasks) and Stage 3 multi-task fine-tuning (73.8 percent, 1.91M samples). Stages 2 and 4 are adapter-merging operations that do not consume new training samples and are omitted from the pie. Total samples processed across both training stages: 2.58M. (c) Donut chart of training-data source composition by cell count (approximately 108 million cells): cellxgene (~35 percent), salusSTS in-house 0.1-micrometer subcellular platform (~17 percent, planned), MOSTA/HESTIA embryo (~15 percent), 10x Visium (~12 percent), Stereo-seq (~9 percent), MERFISH (~5 percent), Replogle 2022 Perturb-seq (~4 percent), sci-Plex3 drug screen (~2 percent), STRING v12 PPI (~1 percent). The salusSTS platform provides twenty million mouse-organ tissue cells with bin40 / bin100 / bin400 multi-scale outputs at 0.1-micrometer subcellular resolution.

### Supplementary Figure S2

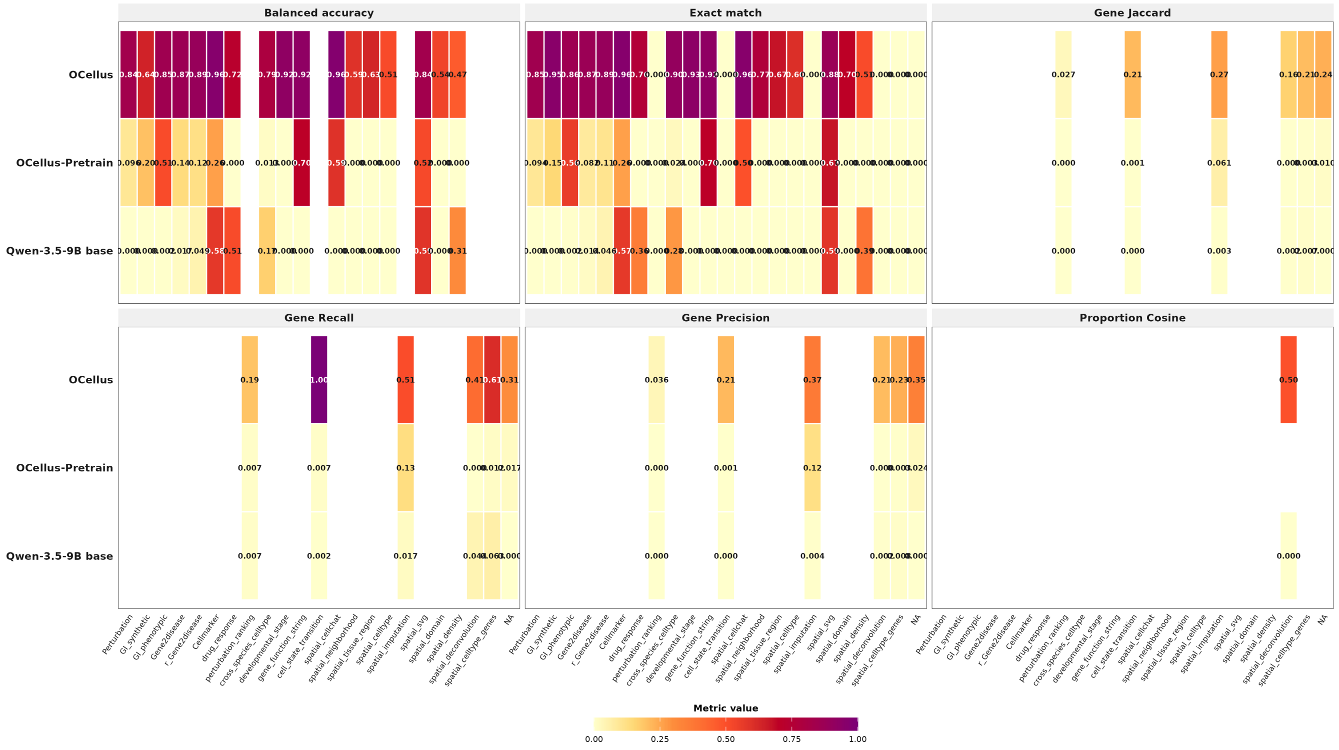


**Supplementary Figure S2. Complete multi-task benchmark heatmap with all six metrics.** Twenty-two evaluated tasks × three model configurations (Qwen-3.5-9B base, OCellus-Pretrain, OCellus) × six metrics: balanced accuracy (mean of per-class recall), exact match (string-level), gene Jaccard (set overlap), gene recall (|A∩B|/|B|), gene precision (|A∩B|/|A|), and proportion cosine similarity (cell-type composition vectors). Each row is one task; each panel is one metric. The primary metric per task (used in Figure 2a of the main text) is highlighted in **bold** in Supplementary Table S6. Empty cells indicate metrics not applicable to that task (e.g., proportion cosine is computed only for spatial deconvolution).

### Supplementary Figure S3

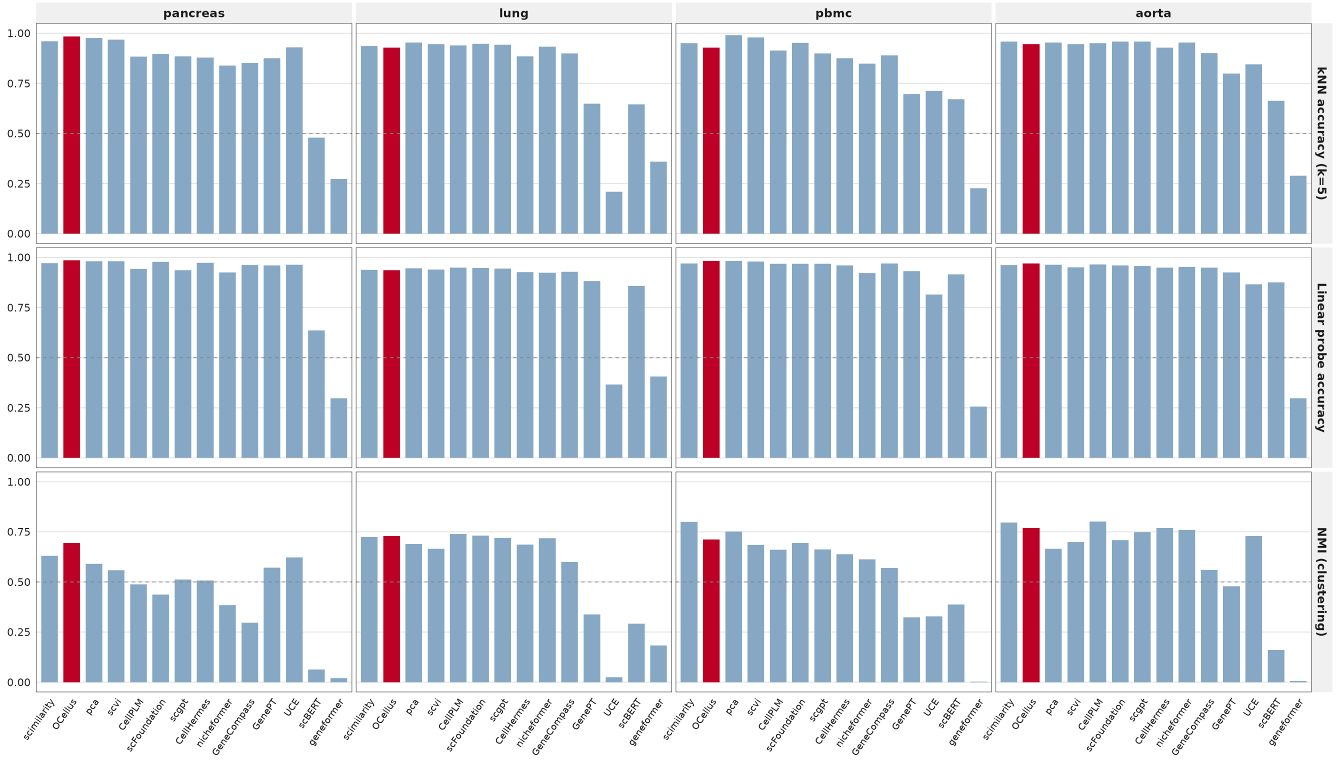


**Supplementary Figure S3. Per-dataset encoder benchmark across 14 foundation models.** For each of the four benchmark datasets (pancreas, lung, PBMC, aorta — top to bottom in each column) we report three metrics (left to right): k-nearest-neighbor accuracy at k=5, linear probe accuracy (logistic regression with five-fold cross-validation), and normalized mutual information against ground-truth cell-type labels. Models are sorted within each panel by metric value. OCellus is highlighted in dark red. Dashed line at 0.5 marks random-chance baseline for kNN and linear probe. The per-dataset breakdown shows that OCellus's encoder advantage is consistent across datasets, though the margin is largest on pancreas and PBMC and smaller on lung and aorta.

#### Supplementary Figure S4

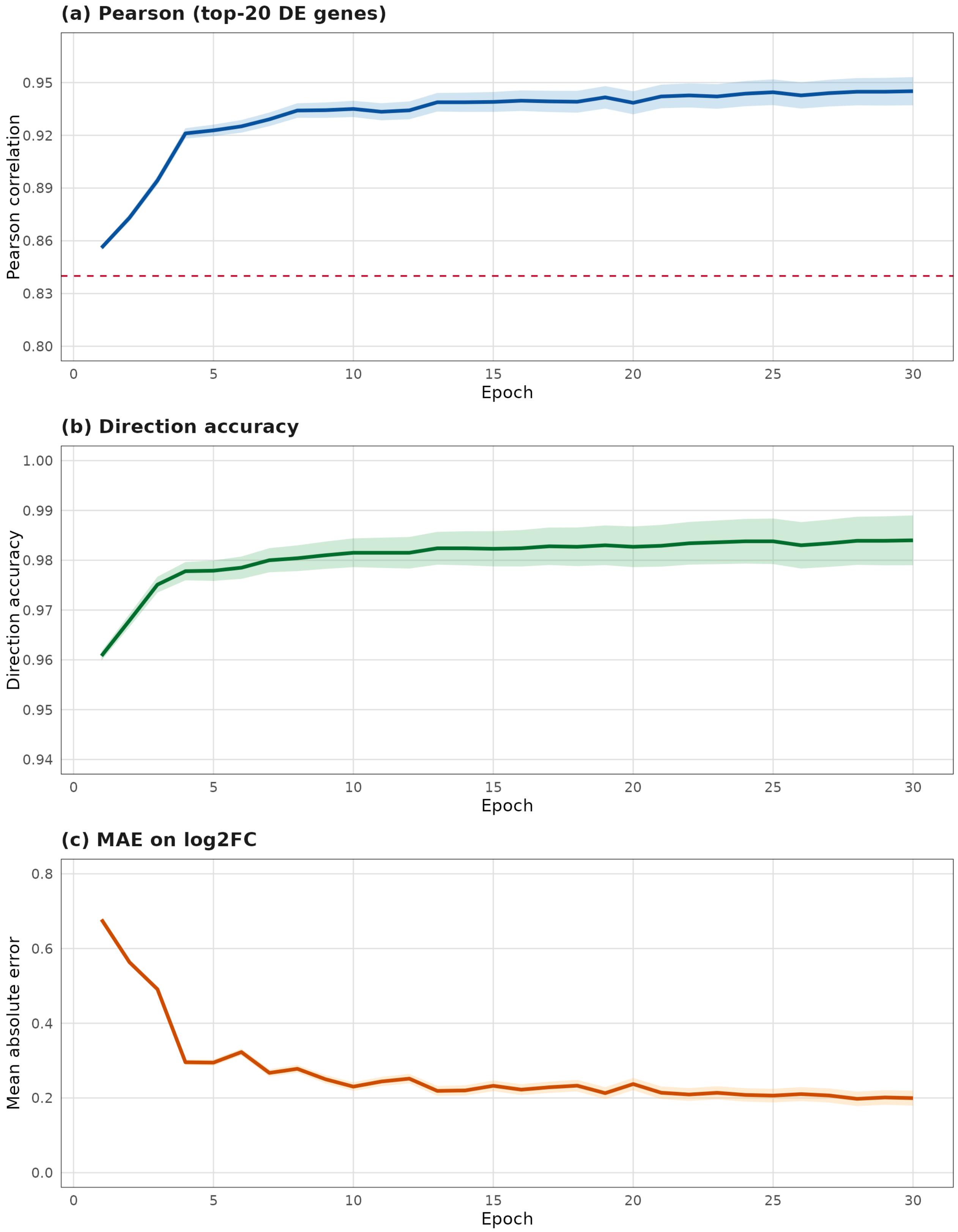


**Supplementary Figure S4. Graph neural network training curves over 30 epochs on the held-out test set.** (a) Pearson correlation on the top-twenty differentially expressed genes per perturbation. The shaded band reflects per-fold variance propagated from the final-epoch three-seed standard deviation (0.008) across the training trajectory; the final reported Pearson is 0.945 ± 0.008. The GEARS baseline (Pearson 0.84, reproduced under our pseudo-bulk top-20-DE evaluation protocol) is marked as a dashed red line. OCellus-GNN surpasses GEARS from epoch 4 onward and continues to improve monotonically through epoch 30. (b) Direction accuracy (binary up/down agreement on the predicted-response-gene ∩ true-DE subset). Final value 98.4 percent; random baseline 0.50. (c) Mean absolute error on log2 fold-change values; final MAE 0.20.
