## Supplementary Tables for "OCellus: A Language-Model Framework for Single-Cell, Spatial, and Perturbation Biology with Natural-Language Reasoning"

### OCellus Paper — Main Tables with Legends

*All 4 main tables with captions.*

#### Table 1

Representative OCellus predictions across six task categories (all six predictions match ground truth; inputs abbreviated for display).

| Task category | Input (abbreviated) | Ground truth | OCellus prediction |
|---|---|---|---|
| Spatial cell type | Atp5j, Uqcrh, Ndufa3, Atp5g3, Gnas, Cox6a1, Mcl1, Atp5b, Ezr, ... | Choroid Plexus | Choroid Plexus ✓ |
| Perturbation (SC) | Gene A = CARF (Activation), Gene B = XPO6, Embryonic stem cells | Yes (Up) | Yes (Up) ✓ |
| Cross-species cell type | Mouse; Slc17a7, Plp1, Rxfp1, Satb2, Tbr1, Sptan1, Mbp, Grin2b, Myt1l, ... | ExN | ExN ✓ |
| Developmental stage | Human; jaw/tooth; ST6GALNAC3, NONO, VIM, SERF2, SPOCK3, VCAN, CADM1, ... | CS18 | CS18 ✓ |
| Synthetic lethality (GI) | Gene A = ETAA1, Gene B = ZNF609 | Synthetic Lethality | Synthetic Lethality ✓ |
| Cell marker identification | Markers MPZ, PAX3, PLP1, POSTN, SOX10, TFAP2A; candidates plasma cell / ... | Cranial neural crest cell | Cranial neural crest cell ✓ |

#### Table 2

Perturbation case study — top-5 response genes following TP53 knockout in K562/HeLA. Numerical predictions are generated under LoRA-enabled mode; biological interpretations are generated under LoRA-disabled mode. Literature reference: Mello et al., *Cancer Cell* (2024); Valente et al., *Cell Reports* (2023).

| Response gene | Predicted FC bin | log2FC | Biological interpretation (OCellus explainer mode) |
|---|---|---|---|
| MDM2 | down_2 | –2.5 | Direct transcriptional target of TP53; loss of feedback regulation |
| CDKN1A (p21) | down_2 | –2.1 | Cell-cycle arrest pathway deactivated |
| BAX | down_1 | –1.4 | Pro-apoptotic signaling reduced |
| SERPINE1 | up_2 | +1.8 | Senescence-associated secretory phenotype (SASP) compensatory activation |
| PMAIP1 (NOXA) | down_1 | –1.2 | Intrinsic apoptosis pathway dampened |

#### Table 3

OCellus-Agent-Bench three-track evaluation.

| Track | Metric | Dataset | OCellus-Agent | Baseline |
|---|---|---|---|---|
| Pipeline accuracy | Task success rate (Verifier-L3 ACCEPT) | 20 NL queries × 4 datasets (80 trials) | 75.0% (60/80) | 51.3% (flat ReAct loop) |
| Perturbation | Pearson (top-20 DE genes) | Replogle 2022 K562 (457 unseen KO) | 0.945 | 0.84 (GEARS) |
| Perturbation | Direction accuracy | Same | 98.4% | — |
| NL explanation | Domain-expert Likert (1–5) | 50 predictions, 3 raters | 4.2 ± 0.6 | 3.1 ± 0.8 (Qwen3.5-9B base) |

#### Table 4

Ablation study — OCellus-Agent pipeline accuracy.

| Configuration | Success rate | Δ vs full | What is removed |
|---|---|---|---|
| OCellus-Agent (full) | 75.0% (60/80) | — | — |
| No Layer-3 Critic | 58.8% (47/80) | –16.2 pp | Critic LoRA disabled, only Layers 1–2 verify |
| Flat ReAct loop | 51.3% (41/80) | –23.7 pp | Coordinator DAG removed, single-turn ReAct |
| No Verifier (all layers) | 51.3% (41/80) | –23.7 pp | All three Verifier layers disabled |
| No LoRA switching | 42.5% (34/80) | –32.5 pp | All tasks use base Qwen3.5-9B (no expert LoRAs) |
